# Host Microenvironment Reprogramming by Saccharides Overcomes Lung Barriers for mRNA Therapeutics

**DOI:** 10.1101/2025.07.14.664686

**Authors:** Lifeng Xu, Tingting Chen, Chao Li, Rui Liao, Qin Xiao, Yang Chen, Fating Yang, Mingxing Luo, Ming Zhang, Shan Guan

## Abstract

Overcoming biological barriers remains the paramount challenge for pulmonary mRNA therapeutics. Conventional approaches focus exclusively on passively optimizing formulation quality without controlling dynamic host barriers. Here, we pioneer a host-centric strategy by leveraging sugar that actively reprograms the airway microenvironment to boost IVT-mRNA transfection. Utilizing machine learning-accelerated screening of a chemically diverse saccharide library, we identify D-glucose as the best-performing candidate. Glucose assisted-delivery within lipid nanoparticles (Glu-LNP) achieves robust, lung-specific protein expression (up to 131.21-fold increase) across diverse preclinical models with reduced inflammation. In lung carcinoma models, Glu-LNP-encapsulated IL-12 mRNA reduced tumor burden by approximately 59.12% and improved survival by 2.5-fold compared to the LNP group. Mechanistically, glucose orchestrates a dual-pathway cascade: metabolic reprogramming via the Warburg effect elevates ATP, fueling endocytosis and translation; ATP further activates the P2Y2-IP3 signaling axis that triggers Ca2^+^ release and subsequent CLCA1/TMEM16A-dependent chloride/bicarbonate efflux, which remodels mucus barriers and enhances nanoparticle penetration. This bioenergetic and mucolytic host intervention strategy presents a broadly applicable paradigm to transcend delivery limitations for respiratory mRNA therapeutics.

## Introduction

The development of airway-applied in vitro-transcribed messenger RNA (IVT-mRNA) therapeutics is highly appealing yet particularly challenging.^1, 2^ Their clinical success is limited by the sophisticated respiratory system and insurmountable biological barriers, including dense mucus layers, hyper-active mucociliary clearance mechanism, and innate immune recognition^3^. Improving IVT-mRNA transfection is critical to ensure adequate protein translation and its downstream biological effects.^4^ Most strategies focus on optimizing the delivery system, including surface condition adjustments of lipid nanoparticle (LNP), large-scale lipid component screening or integration of mucus-penetrating components.^5–9^ Complementary approaches improve IVT-mRNA components via tailored sequence design and high standard purification.^10^ Despite their potential, these approaches remains fundamentally centered on the IVT-mRNA or its vehicle, with limited influence on the dynamic host-dependent biological barriers. None of the previous studies, to the best of our knowledge, has revealed whether safe and transient reprogramming the host environment to actively attenuate biological barriers could concurrently boost IVT-mRNA transfection.

Recent evidences suggest that cellular bioenergetic states critically regulate endocytosis and protein synthesis-essential steps for IVT-mRNA transfection.^11, 12^ The “Warburg effect”, characterized by elevated aerobic glycolysis to meet acute adenosine triphosphate (ATP) demands in rapidly proliferating cells, is particularly relevant. ^13^Such metabolic shifts may similarly enhance IVT-mRNA transfection by improving energy availability and metabolic fitness. Inspired by transcriptomic and metabolomic data derived from our previously tailor-designed pulmonary delivery system (PoLixNano), we serendipitously observed that PoLixNano treatment markedly activated glycolysis and the tricarboxylic acid (TCA) cycle, and upregulated pathways related to uridine diphosphate (UDP) glucuronic acid biosynthesis, suggesting sugar-associated metabolic reprogramming plays critical roles in mediating IVT-mRNA transfection. These observations prompted us to investigate whether tailored manipulation of host metabolic states—utilizing the “Warburg effect” via sugars—could amplify IVT-mRNA transfection.

To this end, we screened 50 commercially available sugars/derivatives at four concentrations and confirmed their ability to enhance LNP-mediated IVT-mRNA transfection in airway epithelia. To further expand the chemical space and identify best-performing candidates, we employed machine learning (ML)-guided virtual screening based on key physicochemical descriptors of sugars to predict IVT-mRNA transfection efficiency. A eXtreme gradient boosting (XGBOOST) model identified physicochemical profiles for top-performing sugars and accurately predicted their in vivo transfection efficacy. D-glucose emerged as the most potent candidate across in vitro and in vivo validations. Notably, D-glucose enhanced IVT-mRNA transfection across diverse nanocarrier systems, including lipid nanoparticles (LNPs), polymeric nanoparticles (PolixNano), and PBAE-based formulations. The optimized D-glucose containing LNP (Glu-LNP) achieved 6.39-fold higher IVT-mRNA expression in cultured cells, 131.21-fold improvement in animal models. Notably, D-glucose demonstrated superior biocompatibility and significantly reduced pulmonary inflammation in rodents. In a melanoma lung metastasis model, Glu-LNP enabled IVT-mRNA encoding interleukin-12 (IL-12) exhibit superior therapeutic outcomes compared to conventional LNP. Tumor burden, as measured by bioluminescence signal, was reduced by 59.12%, survival rate improved by 2.5-fold, demonstrating therapeutic potential via host-directed modulation.

Mechanistic studies revealed D-glucose enhances IVT-mRNA transfection through dual pathway involving metabolic reprogramming via the “Warburg effect” and activation of the P2Y2-IP₃-CLCA1/TMEM16A signalling axis that effectively modulates the mucus barriers.^14–17^ D-glucose elevates intracellular ATP levels by promoting aerobic glycolysis (i.e. the “Warburg effect”), thereby energizing efficient endocytosis and protein translation. Simultaneously, the increased ATP activates purinergic receptor P2Y2, triggering phospholipase C-mediated hydrolysis of phosphatidylinositol 4,5-bisphosphate (PIP₂) into inositol triphosphate (IP₃).^18–20^ This in turn induced calcium ion (Ca²⁺) release from the endoplasmic reticulum, which upregulated calcium-activated chloride channel regulator 1 (CLCA1) and the epithelial chloride channel TMEM16A. Resulting efflux of chloride (Cl⁻) and bicarbonate (HCO₃⁻) ions actively the local mucus microenvironment by decreasing its viscosity and enabling robust IVT-mRNA nanoparticle penetration, therefore prominently ensuring successful IVT-mRNA transfection.

## Results and discussion

### ML-guided identification of best-performing sugar candidates that boost IVT-mRNA transfection

Our initial insights into the metabolic impact on IVT-mRNA transfection were gained through integrated transcriptomic and metabolomic analyses on a previously developed formulation named PolixNano. These data revealed broad alterations in host metabolic pathways, particularly in carbohydrate metabolism (**Extended Data** Fig. 1a,1b), with multiple sugar-related pathways (including glucuronate interconversion and pentose phosphate pathways) significantly enriched (**Extended Data** Fig. 1c,1d). Notably, gene expression associated with glycolysis and the tricarboxylic acid (TCA) cycle was markedly upregulated (**Fig.1a**), accompanied by elevated expression of UDP-glucuronic acid (UDP-GlcUA) pathway related genes and corresponding metabolic intermediates (**Fig. 1b)**, indicating enhanced flux through this biosynthetic route. These observations underscore the crucial roles of sugar metabolism in facilitating efficient IVT-mRNA transfection, which suggests the presence of sugar in IVT-mRNA formulations may potentially improve their efficacy.

**Fig. 1.**
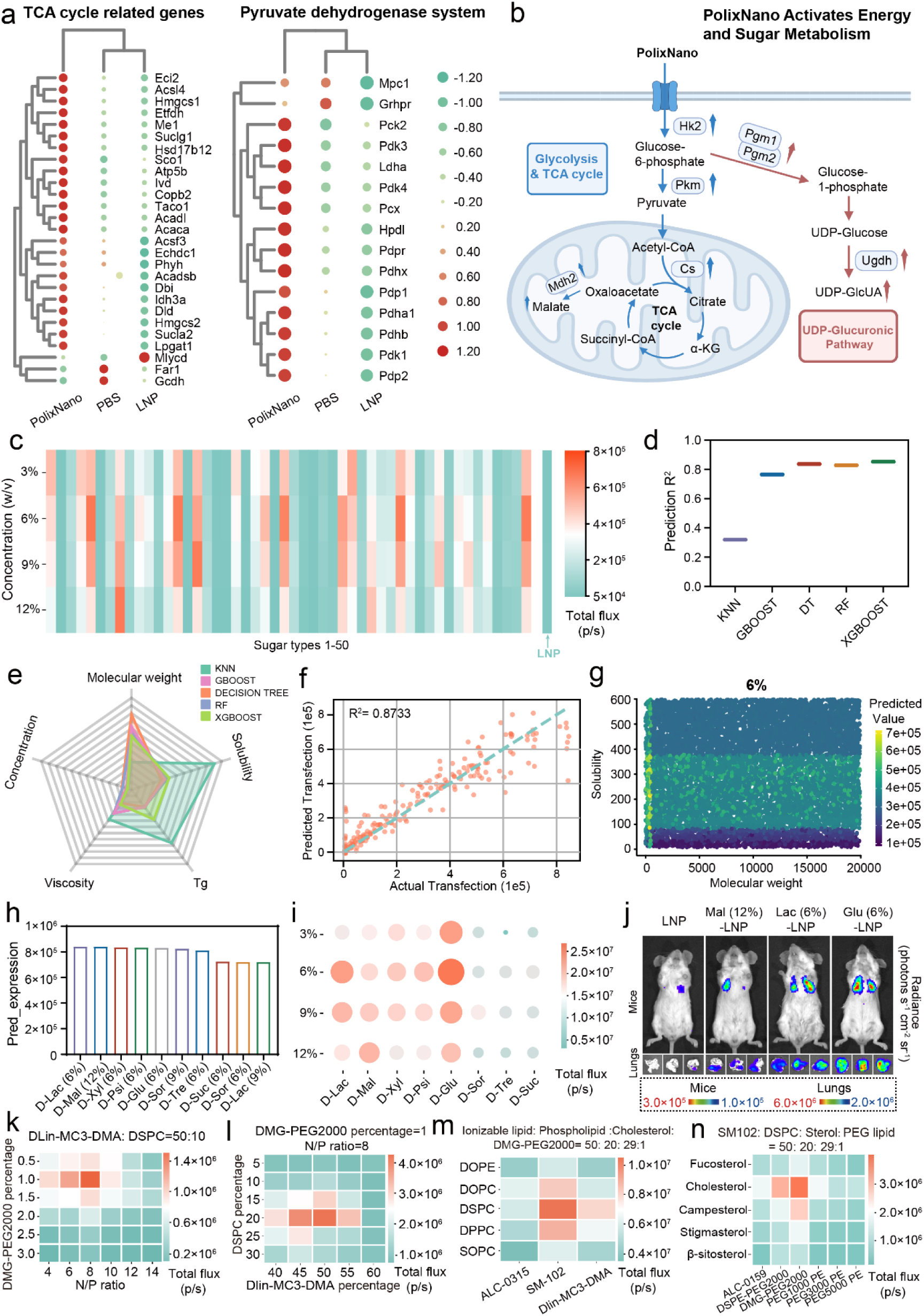
Machine Learning–Guided Sugar Screening for Enhanced LNP-Mediated IVT-mRNA Transfection. **a**, Comparative heatmap analysis of gene expression patterns in the TCA cycle (left panel) and pyruvate dehydrogenase system (right panel) across three experimental groups: PolixNano (an airway tailor-designed mRNA delivery system), LNP and PBS control. Color gradient indicates transcriptional regulation levels (red: upregulated; green: downregulated) with intensity reflecting fold-change magnitude. Gene clustering reflects hierarchical relationships in expression profiles. **b,** Schematic illustration of PolixNano-derived metabolic remodeling related to energy and sugar metabolism. Enzymes with upregulated gene expression are marked alongside their associated metabolic intermediates and mapped onto three main pathways: glycolysis, tricarboxylic acid (TCA) cycle, and the Uridine diphosphoglucuronic acid (UDP-GlcUA) biosynthetic pathway, metabolic flux directionality is indicated by arrows (blue: glycolysis & TCA, red: UDP-GlcUA). Genes are denoted in boxed labels, whereas unboxed elements represent corresponding metabolic intermediates. **c,** Heatmap showing luciferase expression in human bronchial epithelial (16HBE) cells transfected with LNP (containing IVT-mRNA encoding firefly luciferase, i.e. mFluc) co-delivered with 50 different types sugars (sugar-LNP) at four different concentrations (3%, 6%, 9%, and 12% w/v). **d,** Prediction R² values of five different machine learning (ML) models used to evaluate the accuracy of transfection outcome prediction: K-nearest neighbors (KNN), Gradient boosting (GBOOST), Decision tree (DT), Random forest (RF), and Extreme gradient boosting (XGBOOST). **e,** Radar plot showing the importance of solubility, molecular weight, viscosity, and glass transition temperature (Tg) in different ML models for transfection prediction. **f,** Scatter plot showing the relationship between actual and predicted mFluc transfection values for sugar-LNP formulations in the test set, as predicted by the XGBOOST model. Each point represents a single formulation, and the dashed line indicates the linear fit between predicted and actual values. **g–h,** Predicted mFluc transfection values of sugar-LNP formulations screened in silico using a XGBOOST model based on molecular weight and solubility parameters (g), and top 10 best-performing commercially available sugars along with their optimal concentrations (w/v) as ranked by model-predicted scores (h). The selected sugars include D-Lactitol (D-Lac), D-Maltitol (D-Mal), D-Xylitol (D-Xyl), D-Psicose (D-Psi), D-Glucose (D-Glu), D-Sorbitol (D-Sor), D-Trehalose (D-Tre), D-Sucrose (D-Suc). **i,** In vivo mFluc transfection efficiency of the selected 8 sugar-LNP formulations (from h), evaluated in mouse lungs at four concentrations via intranasal administration. Luciferase expression was quantified 6 h post-administration using bioluminescence imaging (n=3 biologically independent animals). **j,** Representative bioluminescence images of mFluc expression in mice and excised lungs 6 h post intranasal administration mediated by LNP formulations supplemented with the top 3-performing sugars identified in (i), namely D-Glu (6% w/v), D-Lac (6% w/v), and D-Mal (12% w/v). Corresponding LNP formulations without sugar served as controls. (n=3 biologically independent animals). **k-n,** Heatmaps illustrating the mFluc transfection efficiency in 16HBE cells mediated by LNP formulations containing 6% (w/v) D-Glucose (Glu-LNP) with different lipid compositions.

To explore this hypothesis, we first validated the transfection-enhancing potential of sucrose, a commonly used lyoprotectant in pharmaceutical compositions. We prepared LNP with D-Sucrose (Suc-LNP) at various concentrations (1%, 3%, 6%, 9%, 12%, and 15% w/v) and evaluated their transfection efficiency in 16HBE cells using IVT-mRNA encoding firefly luciferase (mFluc). Compared to LNP without sucrose, all tested concentrations of Suc-LNP demonstrated significantly improved mFluc expression **(Extended Data Fig.1e**). Encouraged by these results, we further screened 50 types of commercially available sugars and their derivatives at different concentrations (200 distinct formulations) with a simple addition into LNP encapsulating mFluc, and their transfection efficiency in 16HBE cells were subsequently evaluated. The results suggest that most of these sugar-containing formulations mediates significantly higher transfection than the conventional LNP counterpart (**Fig.1c**).

In order to rapidly identify the best-performing sugar candidates for in vivo settings, we applied machine learning (ML) techniques that demonstrates considerable promise in delivery system design and chemical compound screening.^21–25^ In contrast to conventional high-throughput screening, which is often labor-intensive, repetitive, and costly, ML-driven screening enables lower-cost and faster in vivo formulation discovery while substantially reducing the number of animals required for experimental screening.^26–28^ We created an experimental training dataset based on the in vitro transfection profiles of 200 distinct formulations established in the initial screening, and developed computational models to predict sugar-enhanced in vivo transfection efficiency based on their physicochemical features. Specifically, we extracted features of sugars that could be associated with IVT-mRNA transfection, including Molecular weight, Solubility, Viscosity and Glass Transition Temperature (Tg). We found the eXtreme Gradient Boosting (XGBoost) showed the best prediction among five different models, including K-nearest neighbors (KNN), Gradient boosting (GBOOST), Decision tree (DT), Random forest (RF) and XGBoost (**Fig.1d**).

Across all tested models, solubility and molecular weight were consistently highlighted as the most influential features, identifying them as key physicochemical parameters of sugars contributing to IVT-mRNA transfection efficiency (**Fig.1e**). Notably, the correlation between predicted and actual transfection efficiencies validated our models’ accuracy, which could explain 87.3% of transfection variation in the test set (**Fig.1f and Extended Data Fig.2a-c**). Leveraging an XGBoost-based virtual screening of 40,000 simulated sugar-LNP candidates across four different concentrations, we confirmed critical physicochemical ranges (molecular weight: 0-500 Da, solubility: 10-300 mg/mL) that associated with optimal IVT-mRNA transfection (**Extended Data Fig.2a-g and Fig.1g**). Based on GlyTouCan and PubChem databases and commercially available sugars,^29, 30^ we identified the top 10 sugar-concentration formulations with the highest predicted transfection efficiencies according to the selected parameter range (**Fig. 1h**). Subsequent in vivo validation confirmed that all of these top-ranked formulations significantly enhanced LNP-mediated transfection efficiency in the lungs (**Fig. 1i**). Notably, D-Glucose (6% w/v), D-Lactitol (6% w/v), and D-Maltitol (12% w/v) exhibited the highest performance among the tested combinations (**Fig. 1j**). Collectively, these results highlight the potential of sugar-enhanced IVT-mRNA transfection, with D-Glucose at 6% (w/v) emerging as the best-performing candidate.

After identifying the lead sugar candidate, we utilized orthogonal design of experiments (DOE) to systematically optimize lipid compositions for Glu-LNP. Initial analyses evaluated N/P ratios (4-14) and DMG-PEG2000 molar ratios (0.5-3%), with a fixed DLin-MC3-DMA:DSPC ratio at 50:10 (**Fig.1k**). Further optimizations examined varying DSPC (5-30%) and DLin-MC3-DMA (40-60%) molar ratios with a fixed 1% DMG-PEG2000 (**Fig.1l**). Final refinement involved screening diverse types of helper phospholipids, ionizable lipids, sterol components and PEG-lipids (**Fig.1m,1n**), which turns out Glu-LNP formulated with SM-102, DSPC, cholesterol, and DMG-PEG2000 (50%: 20%: 29%: 1% molar ratio, N/P=8) being the optimal lipid composition in presence of D-Glucose.

We next evaluated the physicochemical properties and storage stability of Glu-LNP. Compared to conventional LNP, Glu-LNP exhibited similar particle size, PDI, encapsulation efficiency, viscosity, and maintained a comparable spherical morphology (**Extended Data** Fig. 3a-c). In contrast, Glu-LNP demonstrated improved stability across various temperature conditions (−80 °C, 4 °C, room temperature, and 37 °C), maintaining better physicochemical characteristics over time (**Extended Data** Fig. 3d). Collectively, the ML-guided screening established a highly effective paradigm of sugar boosted IVT-mRNA transfection, with enhanced storage stability of LNP-based formulations.

### The glucose enhances cellular uptake of IVT-mRNA via modulating endocytic mechanisms and promotes in vitro mRNA transfection

To evaluate the impact of glucose on cellular internalization of LNP, we encapsulated Fluorescein-labelled mRNA (mFITC) into Glu-LNP or LNP and analyzed their uptake in 16HBE, DC2.4, and A549 cells. Flow cytometry suggests glucose markedly increased cellular uptake of Glu-LNP across all tested cell types compared to LNP counterparts, as evidenced by elevated mean fluorescence intensity (MFI) and higher percentages of FITC-positive cells (**Fig. 2a**). To further reveal underlying mechanism, a series of endocytosis inhibitors were applied. Compared to LNP control, Glu-LNP treated cells exhibited enhanced sensitivity to both sodium azide and amiloride, indicating that glucose induced energy-dependent and macropinocytosis-mediated internalization (**Fig. 2b and Extended Fig. 4a**), thereby reshaping the cellular uptake landscape to enable more efficient mRNA internalization.^31^

**Fig. 2.**
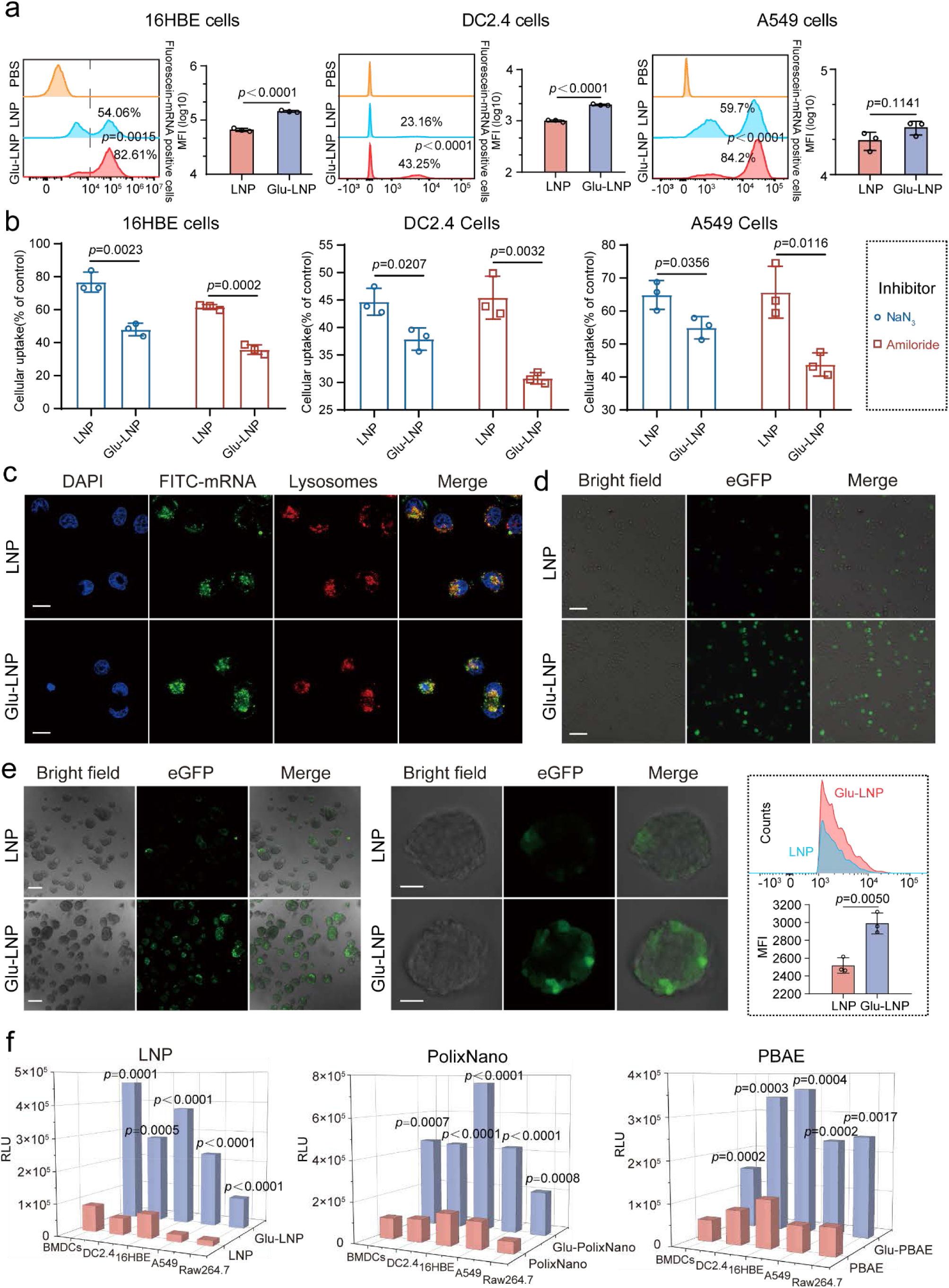
Characterization and in vitro evaluation of glucose co-delivery in enhancing cellular uptake and mRNA transfection efficiency. **a**, The cellular uptake of Fluorescein-mRNA (mFITC) encapsulated in Glu-LNP or LNP counterpart in 16HBE, DC2.4, and A549 cells measured by flow cytometry (n=3 biologically independent samples). The bar chart indicates the mean fluorescence intensity (MFI) per cell, and histograms display the percentage of mFITC positive cells. Untreated cells were used as the blank control. **b,** Mechanistic exploration of cellular uptake pathways using specific endocytosis inhibitors in 16HBE, DC2.4, and A549 cells treated with mFITC encapsulated in Glu-LNP. Control indicates uninhibited uptake (n=3 biologically independent samples). **c,** Subcellular location of the mFITC@Glu-LNP and mFITC@LNP after 6 h incubation with 16HBE cells. Blue channel, nuclei stained by DAPI; green channel, mFITC; red channel, endocytic vesicles marked by LysoTracker™ Red DND-99; Merge, combination of the aforementioned channels. Scale bars: 15 µm. **d,** Representative fluorescence microscopy images of 16HBE cells transfected with Enhanced Green Fluorescent Protein-mRNA (mEGFP) encapsulated in Glu-LNP or LNP after 6 h incubation. Scale bar: 100 µm. **e,** Representative fluorescence microscopy images of organoids transfected with mEGFP encapsulated in Glu-LNP or LNP after 6 h incubation. Images from left to right show a low-magnification bright-field image, EGFP fluorescence (green channel), and the merged overlay. Scale bars: 35 µm (left), 50 µm (middle). **f,** mFluc transfection efficiency across various cell types (BMDCs, DC2.4, 16HBE, A549, RAW264.7) mediated by LNP, PolixNano, PBAE, and their glucose co-delivery counterparts (Glu-LNP, Glu-PolixNano, Glu-PBAE). Results expressed as relative light units (RLU) (n=3 biologically independent samples).

Subcellular localization studies showed that mFITC delivered by either Glu-LNP or LNP was similarly distributed with partial co-localization in lysosomes, indicating that the addition of glucose did not significantly alter lysosomal escape capability of LNP based formulations in 16HBE or DC2.4 cells (**Fig. 2c and Extended Fig. 4b**). However, glucose prominently enhanced the transfection efficiency of mEGFP in 16HBE and DC2.4 cells (**Fig. 2d and Extended Fig. 4c**), and three-dimensional (3D) organoid models. both confocal microscopy and flow cytometry revealed substantially improved ratios of EGFP positive cells in organoids incubated with mEGFP@Glu-LNP (10.57%) compared to mEGFP@LNP (5.46%) (**Fig. 2e**). Besides, glucose barely displayed observable cytotoxicity in all examined cell types (**Extended Fig. 4d**).

To assess the applicability of the glucose mediated Fluc-mRNA transfection enhancement in different types of delivery platforms, LNP (representative lipid-based delivery system), PoLixNano (a polymer-lipid hybrid delivery system) and PBAE (a cationic polymer-based delivery system) were tested in a panel of cell lines (BMDCs, DC2.4, 16HBE, A549, and RAW264.7 cells). Glucose consistently improved the mFluc expression across all delivery platforms tested, highlighting the generalizability of glucose in promoting IVT-mRNA transfection (**Fig. 2f**).

### The glucose enables superior pulmonary IVT-mRNA transfection with better safety profiles

To evaluate the impact of glucose towards pulmonary IVT-mRNA delivery mediated by LNP, we first investigated the biodistribution of fluorescence (DID)-labelled formulations in mice after intranasal administration. The ex vivo imaging reveals that the fluorescence labelled formulation was predominantly localized in the lungs 6 h post-administration, while signal in other major organs was negligible. Quantitative analysis further confirmed that Glu-LNP exhibited significantly higher pulmonary accumulation compared to the LNP counterpart (**Fig. 3a**). To assess transfection profiles, mice were treated by mFluc encapsulated in Glu-LNP or LNP via the same route. Bioluminescence imaging revealed stronger luciferase expression in Glu-LNP treated mice, quantitative analysis showed up to a 45.13-fold increase in pulmonary luminescence compared to the LNP group (**Fig. 3b**), highlighting the capacity of glucose to enhance lung-specific mRNA transfection efficiency.

**Fig. 3.**
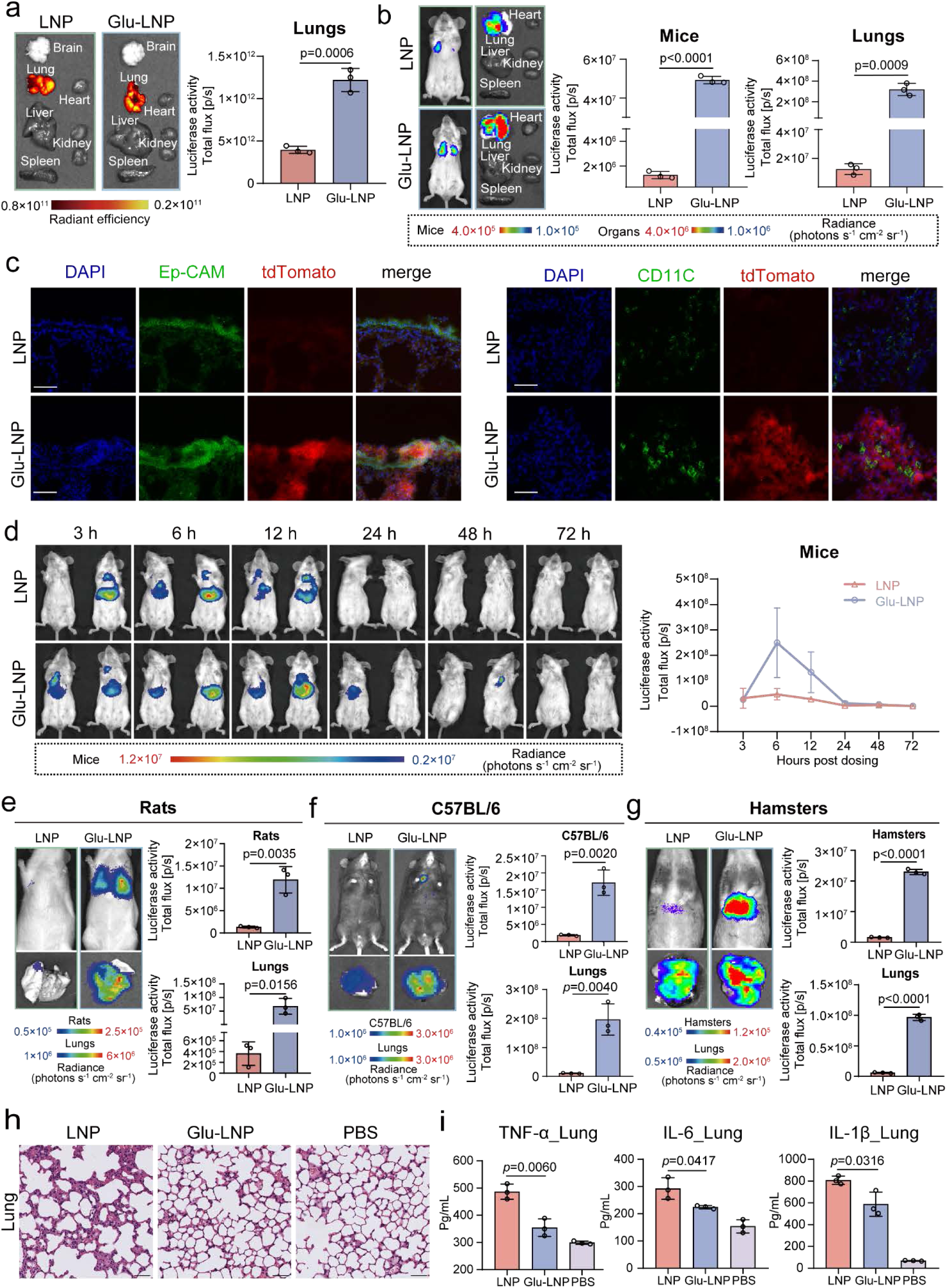
Pulmonary delivery performance and safety profile of Glu-LNP following intranasal administration. **a**, Distribution of fluorescence (DID)-labelled Glu-LNP or LNP in excised organs of mice at 6 h after intranasal instillation (left panel), with corresponding quantitative analysis of fluorescence intensities in lungs (right panel) (n=3 biologically independent animals). **b,** Bioluminescence imaging of mFluc expression mediated by intranasally administered Glu-LNP or LNP in mice and their excised organs at 6 h post-administration (left), alongside quantitative bioluminescence intensities (right) (n=3 biologically independent animals). **c,** Representative immunostaining images illustrating the distribution of Cre-mRNA delivered by Glu-LNP or LNP in lung sections from tdTomato reporter mice at 6 h post-intranasal instillation. For each panel, images from left to right represent nuclei (DAPI, blue), Cre-mediated tdTomato expression (red), EpCAM (epithelial cells, green, left panel) or CD11C (dendritic cells, green, right panel), and the merged image. Scale bar: 50 μm. **d,** Time-course bioluminescence analysis of mFluc expression mediated by intranasally administered Glu-LNP or LNP in mice (n=3 biologically independent animals). **e-g,** Bioluminescence imaging and quantification of mFluc expression in rats, C57BL/6 mice and Hamsters at 6 h post-intranasal instillation of Glu-LNP or LNP (n=3 biologically independent animals). **h,** Hematoxylin and eosin (H&E) staining of lung sections collected from mice treated by Glu-LNP or LNP 6 h post-intranasal dosing. Scale bar: 50 μm. **i,** Concentrations of TNF-α, IL-6, and IL-1β in lung homogenates collected 6 h after intranasal administration of PBS, LNP, or Glu-LNP (n=3 biologically independent samples). Cytokine concentrations were measured by ELISA.

**Fig. 4.**
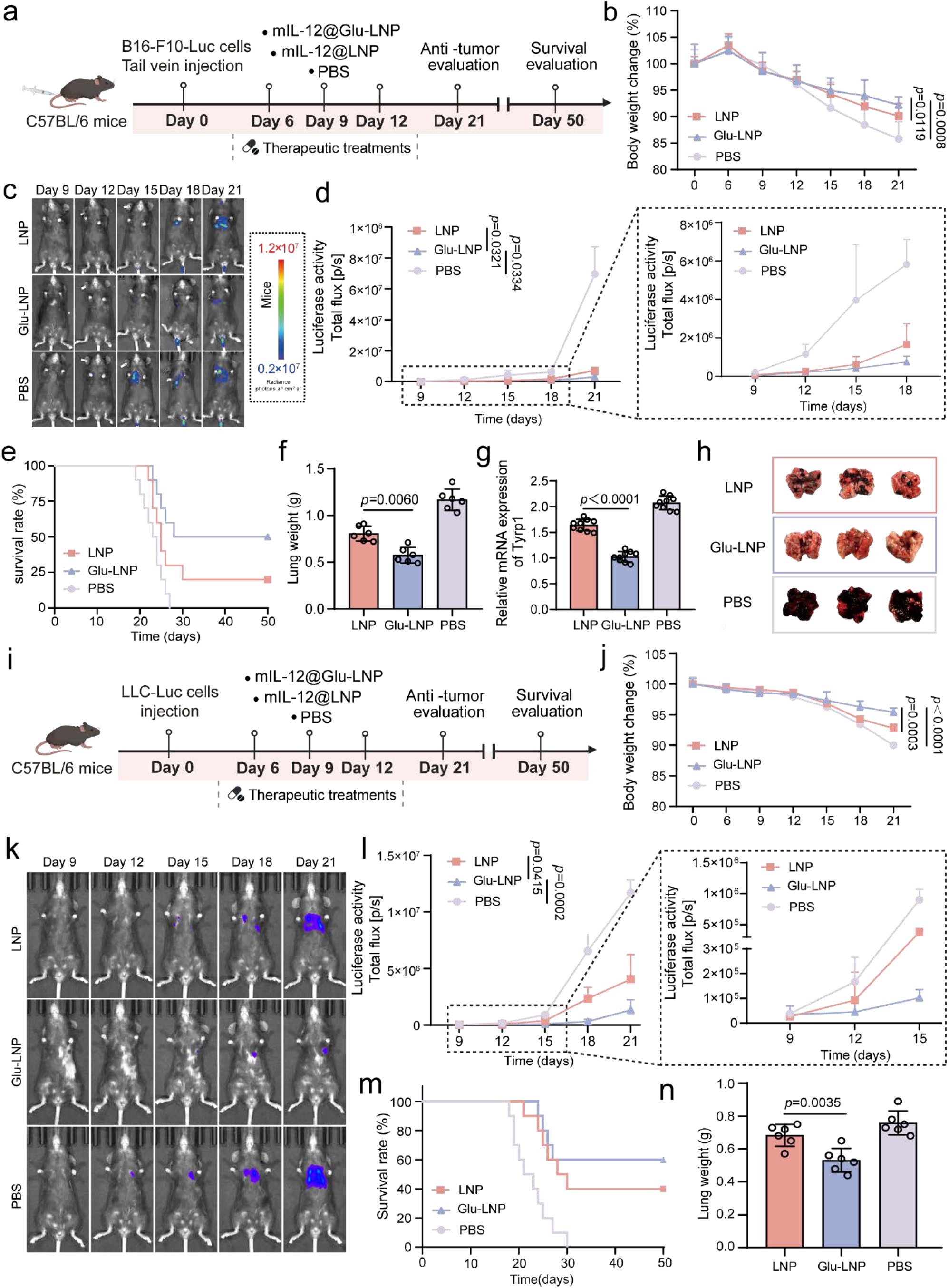
Evaluations of glucose-enhanced therapeutic efficacy of mRNA encoding interleukin-12 (mIL-12) in mouse metastatic and orthotopic lung cancer models. **a**, Schematic illustration of the treatment scheme for the B16-F10-based luciferase expressing melanoma (B16-F10-Luc) induced lung metastasis model in mice. C57BL/6 mice were intranasally administered with PBS, mIL-12@LNP, or mIL-12@Glu-LNP on day 6, 9, and 12 after tumor cell inoculation on day 0. Tumor burden was monitored until day 21, and survival was assessed up to day 50 post-tumor challenge. **b,** Body weight changes of mice treated by the indicated formulations over time (n=9 biologically independent animals). **c-d,** Bioluminescence imaging (c) and corresponding quantification (d) of tumor burden (left panel) on day 9, 12, 15, 18, and 21 (n=3 biologically independent mice). The right panel of (d) represents quantification of luciferase signal intensities at the indicated time points shown in the dashed box in (d), reflecting detailed tumor progression from day 9 to day 18 (n=3 biologically independent animals). Statistical significance was evaluated on day 21 in (b) and (d). **e,** Survival curves of mice treated by indicated formulations (n=9 biologically independent animals). **f,** Lung weight measurement on day 21 (n=6 biologically independent samples). **g,** Reverse Transcription Quantitative Polymerase Chain Reaction (RT-qPCR) analysis on the relative mRNA expression of Tyrosinase-related protein 1 (Tyrp1), a melanoma-associated gene, in lung tissues on day 21 (n=9 biologically independent samples). **h,** Representative images of lungs excised on day 21. Black nodules indicate metastatic foci (n=3 biologically independent mice). **i,** Schematic illustration of the treatment scheme for the Lewis Lung Carcinoma-based luciferase expressing cells (LLC-Luc) established orthotopic lung tumor model. C57BL/6 mice were intranasally administered with PBS, mIL-12@LNP, or mIL-12@Glu-LNP on day 6, 9, and 12 after tumor cell inoculation. Tumor burden was monitored until day 21, and survival was assessed up to day 50 post-tumor challenge. **j,** Body weight changes of mice treated by the indicated formulations over time (n=9 biologically independent animals). **k-l,** Bioluminescence imaging (k) and corresponding quantification (l) of tumor burden (left panel) on day 9, 12, 15, 18, and 21 post-tumor inoculation (n=3 biologically independent animals). The right panel of (l) represents detailed quantification of luciferase signal intensities (reflecting tumor progression) from day 9 to day 18 (n=3 biologically independent animals). Statistical significance was evaluated on day 21 in (j) and (l). **m**, Survival curves of mice treated by PBS, mIL-12@LNP, and mIL-12@Glu-LNP (n=9 biologically independent animals). **n**, Lung weight measurement on day 21 (n=6 biologically independent samples).

To determine Glu-LNP or LNP successfully transfected cell types in the mouse lung, osa26-LSL-tdTomato mice, in which tdTomato expression is induced upon Cre-mediated recombination, were used to trace in vivo mCre (IVT-mRNA encoding Cre) transfection.^32^ Immunofluorescence analysis of lung sections revealed strong tdTomato activation in bronchial epithelial cells (EpCAM+), with a few tdTomato signals being detected in CD11C+ cells in the Glu-LNP group. In contrast, tdTomato signal was barely observable in the LNP-treated counterparts (**Fig. 3c**). The temporal dynamics of glucose enhanced transfection were further assessed by monitoring bioluminescence over 72 h duration. In contrast to a transient and weak signal in the LNP group, Glu-LNP treatment led to stronger and sustained luciferase expression in the lungs, with the peak at 6 h and persistence up to 72 h post-dosing (**Fig. 3d**), indicating favorable expression durability mediated by glucose.

To evaluate the cross-species applicability of glucose-mediated enhancement on IVT-mRNA transfection, Glu-LNP and LNP control formulations were administered intranasally in multiple animal models, including C57BL/6 mice, rats, and hamsters. Glu-LNP effectively mediated mFluc transfection in the lungs of rats, exhibiting a 131.21-fold increase compared to the LNP counterpart (**Fig. 3e**). Similarly, in C57BL/6 mice and hamsters, Glu-LNP demonstrated 26.06-fold and 20.61-fold higher pulmonary mRNA expression, respectively, relative to the LNP controls (**Fig. 3f,3g**). These results demonstrate the consistent and enhanced transfection efficacy of glucose across diverse preclinical models.

To assess the in vivo safety profiles and potential immunomodulatory effects of glucose, histopathological analyses were conducted. Hematoxylin and eosin (H&E) staining of major organs, including the lung, liver, spleen, kidney, heart, and intestine showed no observable pathological alterations in Glu-LNP treated mice compared to LNP and PBS groups, while the lung sections of LNP showed occasionally peribronchial inflammation and neutrophilic infiltration (**Fig. 3h and Extended Fig. 5a**). These observations suggest glucose may alleviate the inflammation stimulating profiles of LNP. To further evaluate local and systemic inflammatory responses, levels of pro-inflammatory cytokines (including TNF-α, IL-6, and IL-1β) in mouse lung tissues, bronchoalveolar lavage fluid (BALF), and serum were measured (**Fig. 3i** and **Extended Fig. 5b**). Compared to LNP treated mice, the Glu-LNP group showed markedly decreased levels of all detected cytokines across all samples, indicating glucose indeed alleviates LNP induced inflammation and improves biocompatibility. In addition, serum biochemical parameters—including ALT, AST, ALP, BUN, CREA, GLU, and GSP remained within normal physiological ranges (**Extended Fig. 5c**), demonstrating safety profiles and physiological compatibility of glucose.

**Fig. 5.**
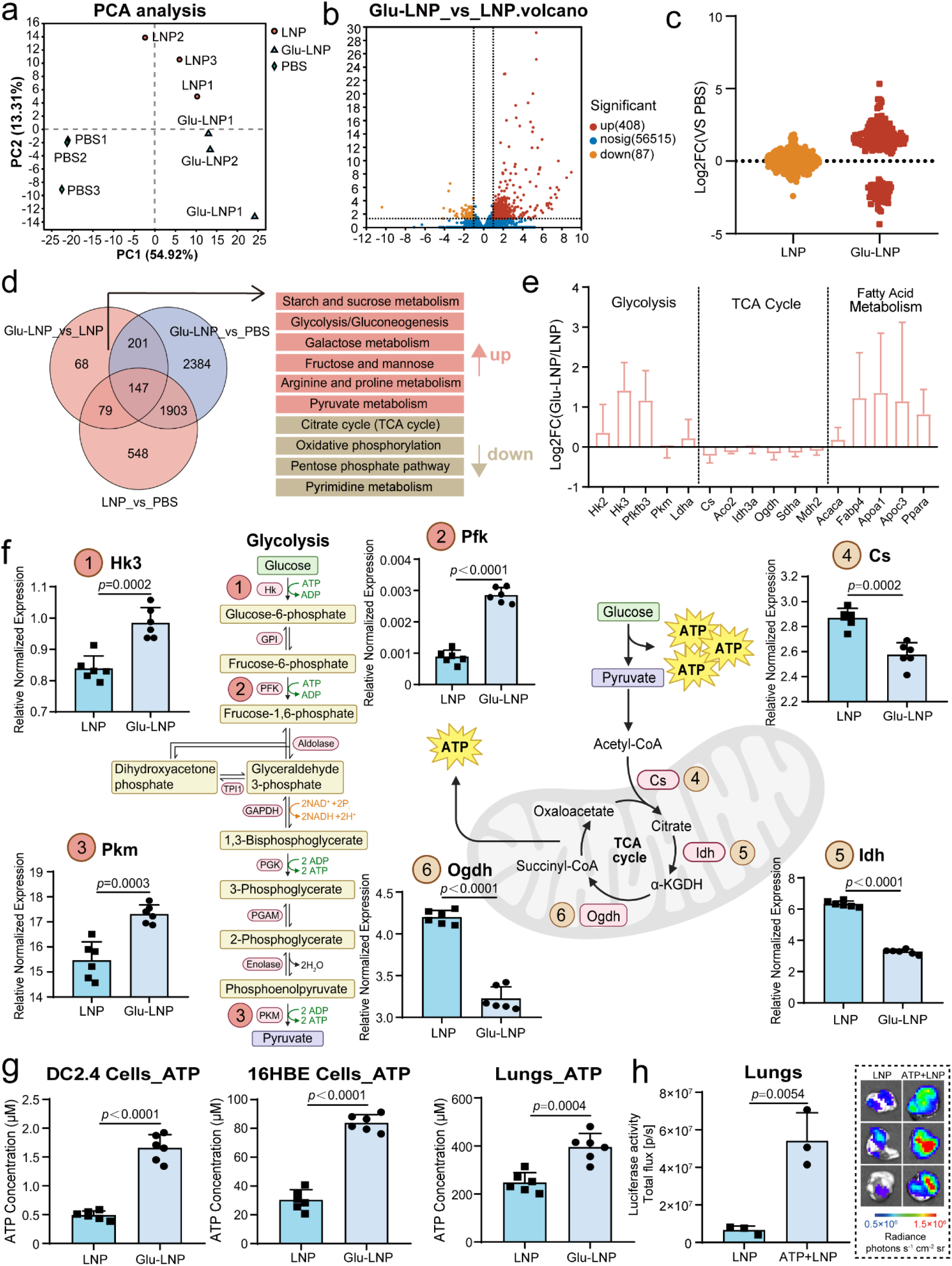
Transcriptomic Analysis reveals Glucose-Driven Metabolic Reprogramming via ATP-Dependent pathways enhancing mRNA Transfection. **a**, Principal component analysis (PCA) of RNA-seq data showing the metabolic profiles of Glu-LNP, LNP, and PBS groups in mouse lung tissue. Each point represents an independent biological sample (n=3 biologically independent mice per group). **b,** Volcano plot of differential gene expression (DEGs) between Glu-LNP and LNP groups. Red dots represent upregulated genes (Log_2_ fold-change, Log_2_FC ≥ 1), and yellow dots represent downregulated genes (Log_2_FC ≤ −1) (n=3 biologically independent samples). **c,** Differentially expressed genes between Glu-LNP and LNP groups relative to the PBS group. The plot shows the Log_2_FC of the significant genes with a *p* adjust value of <0.05 (n=3 biologically independent samples). **d,** Venn diagram illustrating the overlap DEGs between Glu-LNP vs LNP and Glu-LNP vs PBS comparisons. 68 DEGs are identified in Glu-LNP_vs_LNP groups, which are significantly enriched in metabolic pathways and are categorized based on their expression changes that were marked as red (activated) or brown (suppressed). **e,** Log_2_FC of gene expression in key metabolic pathways (glycolysis, TCA cycle, and fatty acid metabolism) between Glu-LNP_vs_LNP groups, based on the 68 DEGs identified in (d). Key genes involved include: (1) Glycolysis: Hexokinase 2(Hk2), Hexokinase 3 (Hk3), 6-phosphofructo-2-kinase/fructose-2,6-biphosphatase 3(Pfkfb3), Pyruvate kinase M (Pkm), Lactate dehydrogenase A (Ldha); (2)TCA Cycle: Citrate Synthase (Cs), Aconitase 2(Aco2), Isocitrate dehydrogenase 3 alpha (Idh3a), Alpha-ketoglutarate dehydrogenase (Ogdh), Succinate dehydrogenase complex subunit A (Sdha), Malate dehydrogenase 2 (Mdh2); (3) Fatty Acid Metabolism: Acetyl-CoA carboxylase (Acaca), Fatty acid synthase (FASN), Fatty acid binding protein 4 (Fabp4), Apolipoprotein A1 (Apoa1), Apolipoprotein C3 (Apoc3), Peroxisome proliferator-activated receptor alpha (Ppara). Log₂FC values represent the magnitude and direction of gene expression changes, with positive values indicating upregulation and negative values indicating downregulation (n=3 biologically independent samples). **f,** Quantitative Real-time polymerase chain reaction (RT-qPCR) analysis showing the relative expression of glycolysis-specific genes (Hk3, Pfk, Pkm) and TCA cycle specific genes (Cs, Idh, Ogdh) in the Glu-LNP and LNP groups. Gene expression data were normalized to *β*-actin as an internal control, and presented as relative expression levels (n=6 biologically independent samples). Glycolysis and TCA cycle pathways were created with BioRender.com. **g,** ATP concentration measurement in 16HBE, DC2.4 cells, and mouse lung tissues 6 h after treatment with Glu-LNP or LNP (n=6 biologically independent samples). **h,** Bioluminescence imaging and quantification of mFluc expression in mouse lung tissues 6 h after administration of LNP or LNP co-delivered with ATP (ATP+LNP). Representative images from fluorescence microscopy are shown on the right (n=3 biologically independent animals).

### Glucose potentiates mRNA therapeutics in murine lung tumor models

To investigate whether glucose could enhance the therapeutic potential of mRNA therapeutics, two murine lung tumor models were established: the B16F10-based luciferase expressing cells (B16-F10-Luc) mediated metastatic lung cancer model and the Lewis Lung Carcinoma-based luciferase expressing cells (LLC-Luc) induced orthotopic lung cancer model. In the B16-F10-Luc model, we encapsulated mRNA encoding interleukin-12 (mIL-12) into Glu-LNP (mIL-12@Glu-LNP) or LNP (mIL-12@LNP),^33, 34^ and intranasally administered to mice according to scheme described in **Fig. 4a**. Mice treated with PBS control and mIL-12@LNP exhibited significant weight loss since Day 12, whereas treatment with mIL-12@Glu-LNP alleviated this trend (**Fig. 4b**). Bioluminescence imaging revealed that mIL-12@Glu-LNP markedly suppressed B16-F10-Luc tumor progression compared to PBS and mIL-12@LNP groups (**Fig. 4c,4d**). Consistently, survival analysis demonstrated a significantly prolonged lifespan (50% survival rate on Day 50) in mIL-12@Glu-LNP group, compared to that only 20% mIL-12@LNP treated mice survived on Day 50 and all PBS-treated mice succumbed on Day 27 (**Fig. 4e**). Lung weights in mIL-12@Glu-LNP group measured on Day 21 were almost halved compared to PBS counterpart (**Fig. 4f**). Correspondingly, expression levels of the melanoma specific marker Tyrosinase-related protein 1 (Tyrp1) were markedly reduced in mIL-12@Glu-LNP group compared with other controls (**Fig. 4g**), correlating with fewer visible metastatic nodules on excised lungs (**Fig. 4h).**

Glucose conferred therapeutic benefits were further validated in an orthotopic LLC-Luc lung tumor model (**Fig. 4i**). The mIL-12@Glu-LNP treatment effectively mitigated weight loss compared to PBS and mIL-12@LNP counterparts (**Fig. 4j**). Bioluminescence imaging confirmed a similar trend on the suppression of tumor growth (**Fig. 4k,4l**). Survival analysis reinforced these findings, showing an improved survival rate of 60% in the mIL-12@Glu-LNP group compared to 40% in the mIL-12@LNP group on Day 50, whereas PBS-treated mice exhibited early mortality (**Fig. 4m**). Lung weights in mIL-12@Glu-LNP group were substantially lower than other counterparts (**Fig. 4n**), consistent with the observed reduction in tumor burden. Together, these findings demonstrate that glucose markedly boosted therapeutic potential of mIL-12 in both metastatic and orthotopic lung tumor models. Improved tumor suppression, prolonged survival, and reduced lung tumor burden collectively support the therapeutic advantage of glucose-containing formulations, underscoring their translational potential for lung cancer treatment.

### Glucose Induced Metabolic Reprogramming Enhances IVT-mRNA Transfection through ATP Production

To investigate underlying mechanisms in which glucose boosts IVT-mRNA transfection, we collected mouse lung samples (treated by Glu-LNP, LNP, and PBS, respectively) and performed a transcriptomic sequencing. The principal component analysis (PCA) revealed clear separation between the PBS, LNP, and Glu-LNP groups (**Fig. 5a**). 408 upregulated and 87 downregulated genes were observed in the Glu-LNP group compared to the LNP group (**Fig. 5b**). The Log_2_ fold change (Log_2_FC) of total differential gene expression (DEGs) revealed the Glu-LNP group exhibited a broader regulation of gene expression compared to the LNP group when normalized to the PBS group (**Fig. 5c**). Venn was performed to eliminate background differences, and identified 68 specifically DEGs in Glu-LNP_vs_LNP groups. Pathway enrichment showed these genes were enriched in metabolic pathways. Glycolysis and fatty acid metabolism were predominantly upregulated, while genes associated with the TCA cycle were downregulated (**Fig. 5d,5e**). Quantitative real-time polymerase chain reaction (qPCR) analysis confirmed significant upregulation of glycolysis-specific genes (Hk3, Pfk, Pkm) and downregulation of TCA cycle-specific genes (Cs, Idh, Ogdh) in Glu-LNP groups compared to LNP counterparts (**Fig. 5f**). This observed genetic changes in metabolic gene expression profile recapitulates the “Warburg effect”, a phenomenon where cells preferentially rely on glycolysis for energy production, even in the presence of oxygen. This metabolic reprogramming suggests the presence of glucose within IVT-mRNA formulations amplifies glycolytic flux, yielding rapid ATP production while simultaneously suppressing the TCA cycle. Such metabolic shift towards glycolysis over oxidative phosphorylation may provide an energetic advantage for supporting resource-intensive processes like IVT-mRNA transfection, where accelerated ATP generation meets the acute bioenergetic demands of nanoparticle delivery and protein synthesis.^35, 36^

To validate our hypothesis, we measured ATP levels in Glu-LNP or LNP treated cells and mouse lungs. Results showed significantly higher ATP levels in Glu-LNP group compared to LNP counterpart (**Fig. 5g**), suggesting that glucose-enriched formulations did enhance host ATP production, which likely contributes to subsequent mRNA transfection. To further confirm the impact of increased ATP on IVT-mRNA transfection, bioluminescence imaging revealed that ATP co-administration (ATP+LNP group) drastically boosted mFluc expression in mice relative to LNP alone (**Fig. 5h**). These results indicate that glucose treatment yields metabolic shift to support more ATP generation, thereby inducing more energy production to support IVT-mRNA transfection.

### Glucose driven CLCA1 activation through P2Y2-IP3-CLCA1 signalling axis impairs mucus barriers via ion flux for enhanced IVT-mRNA transfection

After revealing glucose significantly enhanced glycolysis and ATP production through metabolic reprogramming, we further explored downstream mechanisms by which elevated ATP promotes airway IVT-mRNA transfection. Previous studies suggest ATP can trigger the P2Y2 receptor, initiating intracellular calcium mobilization and subsequent activation of calcium-activated chloride channel regulator 1 (CLCA1), which is critical in modulating chloride ion (Cl^-^) transport and mucus secretion.^37^ Secreted CLCA1, known for its role as a mucus “modulator” facilitates Cl^-^ efflux, thus regulating mucus viscosity and airway hydration.^38, 39^ Indeed, aberrant CLCA1 activity has been associated with impaired Cl^-^ ions and bicarbonate ions (HCO₃⁻) transport,^40^ resulting in increased mucus viscosity and airway obstruction, as observed in cystic fibrosis (CF) patients due to CFTR mutations and decreased CLCA1 expression.^38, 41^ Conversely, enhancing CLCA1 expression significantly ameliorates mucus-related symptoms in CF models.^42^ Given these findings, we hypothesized that glucose-induced enhancement of ATP production might activate a P2Y2/CLCA1 signalling axis in the airway, thereby weakening the host mucus barriers, which in turn improves the mobility and transfection efficiency of IVT-mRNA@LNP nanoparticles.

To this end, the expression patterns of CLCA1-related genes between Glu-LNP and LNP treated samples were analyzed through RNA-seq and qPCR analyses. We observed upregulation of *Slc2a1* (encoding the glucose transporter GLUT1, responsible for cellular glucose uptake), *Hif1a* (hypoxia-inducible factor 1*α*, a master regulator of glycolytic gene expression), *P2ry2* (purinergic receptor P2Y2, a G protein– coupled receptor activated by ATP/UTP that triggers downstream calcium signalling), *Plcb1*, *Plcb2*, *Plcb4* (plcb family encodes phospholipase C beta isoforms, which hydrolyze phosphatidylinositol 4,5-bisphosphate (PIP₂) to generate inositol 1,4,5-trisphosphate (IP₃) and diacylglycerol (DAG), key second messengers for calcium mobilization), *Itpr1* (inositol 1,4,5-trisphosphate receptor type 1, mediating calcium release from the endoplasmic reticulum), *Clca1* and *Ano1* (anoctamin 1, a calcium-activated chloride channel) genes. In contrast, *P2rx7* (purinergic receptor P2X7, an ATP-gated ion channel implicated in inflammatory responses) gene was significantly downregulated in the Glu-LNP group (**Fig. 6a and Extended Fig. 6a**). Western blot analysis further validated the protein expression of these genes, including GLUT1 (encoded by *Slc2a1*), PLCβ1 (encoded by *Plcb1*), IP3R1 (encoded by *Itpr1*), P2RY2 (encoded by *P2ry2*), P2RX7 (encoded by *P2rx7*) and CLCA1 (encoded by *Clca1*), revealing their protein expression trends were consistent with the gene expression patterns as described above (**Fig. 6b and Extended Fig. 6b**). These findings suggest that the glucose induces a shift in host gene expression and activates the P2Y2-IP3R-CLCA1 signalling pathways, which may contribute to the CLCA1-mediated Cl^-^ efflux.

**Fig. 6.**
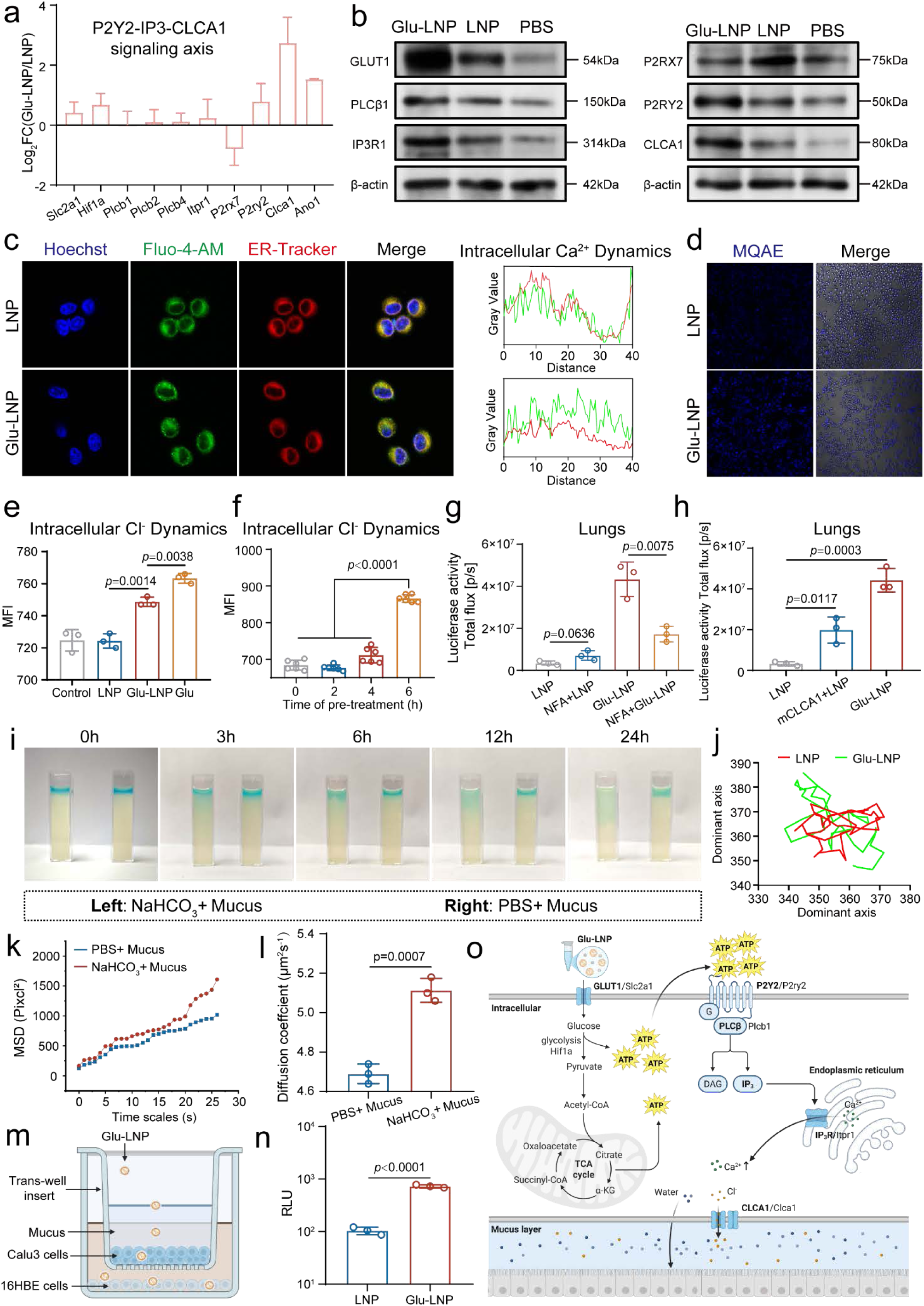
Glucose Activates the P2Y2-IP3-CLCA1 Signalling Axis to Regulate Calcium-Chloride Dynamics and impair mucus barriers. **a**, Differential gene expression analysis showing the expression of key genes associated with the P2Y2 (Purinergic receptor P2Y2)-IP3 (Inositol 1,4,5-triphosphate)-CLCA1 (calcium-activated chloride channel 1) signalling axis. The bar chart in upper left panel represents the Log_2_FC of differentially expressed genes between the Glu-LNP and LNP groups, where a positive value indicates upregulation and a negative value indicates downregulation (n=3 biologically independent samples). **b,** Western blot (WB) analysis showing the protein levels of Glucose transporter 1 (GLUT1), Phospholipase C beta 1 (PLCβ1), Inositol 1,4,5-triphosphate receptor type 1 (IP3R1), Purinergic receptor P2X7 (P2RX7), P2RY2 and CLCA1 in mouse lung tissues treated with Glu-LNP, LNP, or PBS. **c,** Fluorescence imaging showing intracellular calcium dynamics in 16HBE cells treated by Glu-LNP, LNP, or PBS. The left panel shows representative fluorescence images with the blue channel representing nuclei stained by DAPI, the green channel indicating calcium (Fluo-4 AM), and the red channel showing the endoplasmic reticulum (ER tracker). The right panel displays the corresponding fluorescence intensity profiles over distance (x-axis), illustrating the movement of calcium from the ER into the cytosol. **d,** Representative fluorescence images showing intracellular chloride ion concentration in 16HBE cells treated with Glu-LNP or LNP, measured using MQAE dye. **e,** Measurement of intracellular chloride ion dynamics in cells treated with Glu-LNP, LNP, or Glu (glucose only) in the presence of MQAE (n=3 biologically independent samples). **f,** The effect of pre-treatment time on intracellular chloride ion dynamics. 16HBE cells were treated with Glu-LNP for various pre-treatment times (0, 2, 4, 6 h) before measuring chloride ion concentrations using MQAE (n=3 biologically independent samples). **g,** Effect of chloride ion channel inhibitor (NFA, Niflumic acid) on Fluc-mRNA transfection in mouse lung tissues (n=3 biologically independent animals). **h,** Rescue experiment using LNP vectors loaded with mRNA endoding CLCA1 (mCLCA1) administered 6 h prior to the standard mFluc-LNP transfection in mice (n=3 biologically independent animals). **i,** Representative images of cuvette-based diffusion assays, illustrating the diffusion process of fluorescence-labelled nanoparticles within mucus treated with sodium bicarbonate (NaHCO₃, left) or PBS (right) at indicated time intervals (0, 3, 6, 12, 24 h). **j,** Representative nanoparticle tracking analysis (NTA) trajectories of nanoparticles diffusing in mucus samples treated with NaHCO₃ or PBS. **k,** Mean squared displacement (MSD) analysis of LNP nanoparticle trajectories in mucus samples treated with NaHCO₃ or PBS. **l,** Diffusion coefficients calculated from MSD data for LNP nanoparticles in mucus samples treated with NaHCO₃ or PBS (n=3 biologically independent samples). **m,** Schematic illustration of the Transwell assay setup utilizing mucus-secreting Calu-3 cells at the apical side of the insert and 16HBE cells at the bottom, employed to evaluate nanoparticle transmucosal penetration efficiency. **n,** Quantification of basolateral luciferase expression in Transwell experiments following treatment with Glu-LNP or LNP (n=3 biologically independent samples). **o,** Schematic illustration of the mechanism by which the activation of P2Y2 (Purinergic receptor P2Y2)-CLCA1 (calcium-activated chloride channel 1, CLCA1) axis facilitates chloride ion efflux, leading to osmotic water flux, and ultimately contributing to mucus dilution. The receptors are indicated in bold, with their corresponding coding genes shown after the slash. Figure created with BioRender.com.

We further examined intracellular calcium influx and Cl^-^ efflux to elucidate the downstream ionic events. Ca²⁺ are important second messengers in cell signal transduction, and mucus regulation is a calcium-dependent biological process-where spontaneous Ca²⁺ release from the endoplasmic reticulum (ER) has been shown to be a key factor in modulating mucus viscosity.^43^ We observed that calcium ions moved away from the ER into the cytosol of 16HBE cells treated by Glu-LNP, while calcium ions were predominant localized within the ER in the LNP counterpart (**Fig. 6c**). Given that CLCA1 is a calcium-activated chloride channel which mediates Cl^-^ efflux upon calcium elevation, we thus employed MQAE, a probe that tends to be quenched by Cl^-^, to investigated intracellular Cl⁻ levels. The Glu-LNP promoted Cl^-^ efflux compared to the LNP counterpart, as evidenced by a lower intracellular Cl^-^ concentration and a corresponding increase in MQAE fluorescence in the Glu-LNP group (**Fig. 6d**). We examined whether glucose alone (Glu) or Glu-LNP together activates chloride channels. Both Glu and Glu-LNP facilitated Cl⁻ efflux, whereas the LNP counterpart alone had no effect. This indicates that glucose itself activates the chloride channel and enhances Cl⁻ transport, independent of the LNP formulation (**Fig. 6e**). To investigate the temporal dynamics of Cl^-^ flux, we pre-treated the cells with Glu-LNP for different time intervals (0-6 h) and found that Cl^-^ efflux significantly increased during 4-6 h post-treatment (**Fig. 6f**), indicating that glucose-mediated Cl^-^ transport is time-dependent. In addition, we assessed influences of Cl^-^ channel inhibition on IVT-mRNA transfection using a CLCA1 inhibitor—niflumic acid (NFA). When Cl^-^ channels were blocked, no negative impact was observed in LNP mediated IVT-mRNA transfection whereas the Glu-LNP counterpart prominently decreased in presence of NFA (**Fig. 6g**). Based on these observations, we pre-treated mice with IVT-mRNA encoding CLCA1 (mCLCA1) encapsulated in LNP 6 h prior to mFluc@LNP administration, and found this pretreatment moderately enhanced the IVT-mRNA transfection efficiency, though not to the level observed in the Glu-LNP group (**Fig. 6h**).

Previous studies indicate that secreted CLCA1 activates calcium-dependent HCO₃^-^ channels, including transmembrane protein 16A (TMEM16A, also known as ANO1). This activation elevates intracellular Ca²⁺, thereby enhancing TMEM16A/ANO1-mediated HCO₃⁻ transport and promoting abundant HCO₃⁻ secretion into the airway lumen.^40^ The resultant alkaline microenvironment promotes loosening and unfolding of mucin into a reticular structure, forming a gradient mucus layer with decreasing density and increasing permeability from the inner to the outer layer.^40^ This structural change facilitates IVT-mRNA@LNP nanoparticle transport across the mucus barrier. To validate this mechanism, qPCR analysis revealed significantly elevated expression of the TMEM16A/ANO1 gene in the Glu-LNP compared to the LNP group (**Extended Fig. 6a**). Meanwhile, markedly increased HCO₃⁻ concentrations and elevated pH values could be detected in BALF samples collected from Glu-LNP treated mice (**Extended Fig. 6c,6d**), confirming the establishment of a mild alkaline microenvironment. To further validate whether bicarbonate ion efflux could reduce mucus viscosity, we simulated an alkaline microenvironment by directly adding sodium bicarbonate (NaHCO₃) to airway mucus solutions in vitro. In one group, sodium bicarbonate (NaHCO₃) solution was directly added to the mucus solution to create a bicarbonate-rich alkaline environment, while the control group received an equal volume of PBS without NaHCO₃. After 6 h of incubation, viscosity measurements showed a significant reduction in mucus viscosity in the NaHCO₃-treated group compared to the PBS control (**Extended Fig. 6e**). To visually assess whether this change facilitates nanoparticle diffusion, a cuvette-based diffusion assay was performed. Fluorescence-labelled IVT-mRNA@LNP exhibited a markedly larger diffusion range in NaHCO₃-treated mucus than in the PBS-treated counterpart (**Fig. 6i**), suggesting improved mucus permeability under mild alkaline conditions.

We further conducted nanoparticle tracking analysis and multiple-particle tracking to quantify IVT-mRNA@LNP mobility within NaHCO₃-treated mucus samples. Both analyses revealed significantly enhanced diffusion velocity and displacement of IVT-mRNA@LNP in the NaHCO₃-treated group (**Fig. 6j-l**), confirming that bicarbonate-induced alkalization alters mucus structure, effectively reduces mucus barriers and significantly promotes IVT-mRNA@LNP diffusion. A Trans-well assay using mucus-secreting Calu-3 cells at the apical side of Trans-well insert further revealed that glucose pretreatment significantly enhanced mFluc@LNP transmucosal penetration, as evidenced by elevated basolateral luciferase expression compared to LNP counterpart (**Fig. 6m,6n**). Together, glucose treatment did demonstrate positive impacts on the mucus penetration and transepithelial transportation of IVT-mRNA@LNP nanoparticles.

Collectively, these results provide evidences that glucose enhances IVT-mRNA transfection by modulating host mucosal barriers via the P2Y2-IP₃-CLCA1 signalling axis. Specifically, increased intracellular ATP activates the purinergic receptor P2Y2, leading to PLCβ1-mediated generation of IP₃. Elevated IP₃ promotes Ca²⁺ release from intracellular ER stores, triggering Cl^-^ ion efflux and HCO₃⁻ secretion. The resulting mild alkaline environment significantly reduces mucus barrier properties, ultimately facilitating nanoparticle uptake and enhancing IVT-mRNA transfection efficiency (**Fig. 6o**). Notably, the activation of the P2Y2-CLCA1 axis observed in our study echoes the mechanism of Diquafosol ophthalmic solution, a clinically approved P2Y2 receptor agonist. Diquafosol has been shown to promote Cl^-^ efflux by activating the CLCA1/TMEM16A channels, thereby improving mucus hydration and ocular surface homeostasis in dry eye patients.^44, 45^ This clinical precedent reinforces our findings that glucose-triggered activation of the P2Y2-IP₃-CLCA1 cascade enhances ion efflux and modulates the mucus microenvironment to facilitate efficient IVT-mRNA transfection.

## Conclusions

Taken together, this study pioneers a host-centric paradigm that utilizes sugar-mediated metabolic reprogramming and host barriers remodelling strategy for efficient IVT-mRNA transfection, moving beyond conventional approaches focused solely on nanoparticle/mRNA-centric optimizations. Through machine learning–guided virtual screening, we identified D-glucose as a top-performing sugar candidate. By orchestrating the “Warburg effect” (elevating ATP via glycolysis to enhance endocytosis/translation) and purinergic ion-flux pathways (reshaping extracellular mucus barriers), our “host intervention” strategy offers a broadly applicable, biocompatible and scalable solution for respiratory IVT-mRNA therapeutics. Considering sugars are readily available, inherently safe, and broadly adaptable, this strategy holds clinical translation potential for pulmonary mRNA therapeutics across cytokine therapies and metastatic cancer intervention.

## Methods

### Reagents

SM-102 (O02010), ALC-0315 (O02008), DLin-MC3-DMA (O02006) and dipalmitoylphosphatidylcholine (DPPC) (S01004) were purchased from Advanced Vehicle Technology L.T.D. Co (Shanghai, China). Other lipids were purchased from Avanti Polar Lipids. 1,1’-dioctadecyl-3,3,3’,3’-tetramethylindodicarbocyanine perchlorate (DiD) (D7757) was purchased from Thermo Fisher Scientific. All other solvents and reagents were obtained at analytical or HPLC grade from Sigma-Aldrich. All commercially available sugars were purchased from Sigma-Aldrich or MedChemExpress (MCE).

### In vitro transcribed mRNA

In vitro transcribed messenger RNA (IVT mRNA) were prepared and purified as previously described.^10^ Briefly, IVT mRNA (the open reading frame (ORF) sequences of IVT mRNA used in this study can be found in Supplementary Table 1) was synthesized by linearized plasmid templates using the T7 High Yield RNA Synthesis Kit (E131-01A, novoprotein) with N1-methylpseudouridine (Glycogene, Wuhan, China) substitution and followed by the addition of a Cap 1 structure using Cap 1 Capping System (M082-01B, novoprotein). The synthesized IVT mRNA was purified using an affinity chromatography-based purification. The concentration and purity of IVT mRNA were measured using a NanoDropTM One (Thermo Fisher Scientific).

### Preparation of LNPs and modified LNP formulations

Unless otherwise specified, LNP was prepared by mixing an aqueous phase with an ethanol phase using a microfluidic device (AceNANOSOME, ACMEI LIFESCIENCE, Shanghai, China) at a volume ratio of 3:1 with a total flow rate of 12 mL/min. Ethanol phase: Lipid components, including ionizable lipid (SM-102, DLin-MC3-DMA or ALC-0315), helper phospholipid (DOPC, DOPE, DSPC, DPPC or SOPC), cholesterol, and PEG-lipid (DMG-PEG2000), were dissolved in ethanol at a molar ratio of 50:20:29:1, or at specific ratios as indicated in certain investigations. Optionally, hydrophobic polymeric components were dissolved in the ethanol phase. Aqueous phase: IVT mRNA was dissolved in citric acid buffer (50 mM, pH = 4). The nitrogen-to-phosphorus (N/P) ratio between IVT mRNA and the ionizable lipid was 8. DiD-labelled nanoparticles (DiD-LNP) were prepared by adding DiD to the ethanol phase at 1 mol% relative to the total lipid content.

The resulting LNP suspension was dialyzed against 1× PBS (pH 7.4) overnight using a dialysis bag with a molecular weight cutoff of 8-14 kDa at 4 °C for 16 h prior to use. Sugar-assisted LNPs (Sugar-LNPs) were prepared by directly adding sterile-filtered sugar powder (e.g., glucose, sucrose, or lactitol) into the dialyzed LNP suspension at the desired dosage (e.g., 5 mg sugar per 50 μL LNP formulation for a final concentration of 10% w/v). The mixture was gently vortexed (800 rpm, 5 min at 4 °C), followed by additional incubation at room temperature with periodic inversion (15 min, every 3 min) to ensure uniform dispersion. If necessary, PBS was added to adjust the final volume to 50 μL per dose. Sugar was considered successfully incorporated when no visible precipitation occurred and the LNP retained stable dispersion characteristics. ATP-modified LNP (ATP+LNP) were prepared by supplementing the aqueous phase with ATP during LNP assembly. Briefly, ATP (A2383, Sigma-Aldrich) was dissolved in nuclease-free water at a final concentration of 0.2 mg/mL and added to the aqueous phase containing IVT mRNA and citric acid buffer (50 mM, pH 4). The resulting ATP-containing aqueous phase was used directly in the microfluidic mixing process to prepare LNPs following standard procedures. Dialysis and downstream processing were performed as described above.

### Physicochemical characterization and IVT-mRNA encapsulation efficiency (EE) analysis

The particle size and zeta potential of LNP/Glu-LNP formulations were determined using a Zetasizer Nano ZS (Malvern Instruments) at 25°C, while their morphology was examined by transmission electron microscopy (TEM, JEM-1400plus, JEOL Ltd., Japan). For TEM imaging, samples were negatively stained with 2% uranyl acetate. The viscosity of the formulations was measured using a HAAKETM MARST viscometer (Thermo Fisher Scientific). To assess IVT-mRNA encapsulation efficiency (EE), LNP/Glu-LNP samples were dispersed in 1×Tris-EDTA (TE) buffer or lysed with 2% Triton X-100, incubated at 37°C for 30 min, and diluted to final IVT-mRNA concentrations ranging from 0.04 µg/mL to 2 µg/mL. RNA standard solutions were prepared at concentrations of 0 µg/mL, 0.04 µg/mL, 0.2 µg/mL, 1 µg/mL, and 2 µg/mL using 1×TE buffer. For fluorescence measurement, 100 µL of each sample or standard was mixed with 100 µL of RiboGreen reagent (1:200 dilution) in a clear-bottom black plate, incubated at room temperature for 3 min, and analyzed using a VarioskanTM LUX microplate reader (Thermo Fisher Scientific) at excitation and emission wavelengths of 480 nm and 520 nm.

### Cellular uptake, internalization mechanism, and transfection efficiency analysis

Fluorescein-labelled IVT-mRNA was prepared using the Label IT® Nucleic Acid Labeling Kit (MIR 3200, Mirus Bio) following the manufacturer’s protocol. For cellular uptake studies, DC2.4 cells (1×105/well), 16HBE cells (1.5×105/well), A549 cells (2×105/well), and bone marrow-derived dendritic cells (BMDCs, 5×105/well) were plated in 24-well plates and cultured overnight. Upon reaching ∼70% confluency, LNP/Glu-LNP formulations encapsulating 200 ng Fluorescein-mRNA were added to each well. After 4 h incubation, cells were washed twice with PBS, and fluorescein-positive cells were quantified via flow cytometry (BD FACSCanto™ II, BD Biosciences).

To investigate the internalization mechanisms, DC2.4 cells (1×105/well), 16HBE cells (1.5×105/well), or BMDCs (5×105/well) were pre-treated with specific inhibitors: sodium azide (0.5 mg/mL, energy depletion), amiloride (0.5 mM, macropinocytosis inhibition), β-cyclodextrin (1 μg/mL, lipid raft disruption), protamine sulfate (1 mM, heparan sulfate competition), or chlorpromazine (10 μg/mL, clathrin-mediated endocytosis inhibition). Cells were incubated with inhibitors at 37°C for 1 h or pre-chilled at 4°C (energy-dependent uptake inhibition). Fluorescein-mRNA@LNP/Glu-LNP (200 ng mRNA/well) was then added, followed by 4 h incubation under corresponding conditions. Untreated cells receiving Fluorescein-mRNA@LNP/Glu-LNP served as the positive control. Relative uptake rates were normalized to the positive control and analyzed by flow cytometry.

For transfection efficiency assessment, 16HBE cells (3.5×105/well), DC2.4 cells (2×105/well), A549 cells (2.5×105/well), and BMDCs (3×105/well) were seeded in 96-well plates (Corning, 3599) 24 h prior to transfection. Serum-free Opti-MEM (170 μL/well) was added, followed by LNP/Glu-LNP formulations containing 400 ng MetLuc-mRNA (30 μL/well, n ≥ 3). After 6 h incubation, media were replaced with complete medium, and cells were cultured for an additional 24 h. Supernatants (50 μL) were mixed with 30 μL luciferase substrate (QUANTI-Luc™, InvivoGen), and luminescence was measured using a Varioskan™ LUX microplate reader (Thermo Fisher Scientific).

### Cell viability assessment

Cell viability was evaluated using the Cell Counting Kit-8 (CCK-8, HY-K0301, MedChemExpress). Briefly, 16HBE cells (3.5×104/well), DC2.4 cells (2×104/well), and A549 cells (2×104/well) were seeded in 96-well plates and cultured for 24 h. Cells were then treated with LNP/Glu-LNP (400 ng IVT-mRNA/well), naked IVT-mRNA (400 ng/well), 10% (w/v) glucose, PBS, or Lipo3000 (Lipofectamine 3000, Thermo Fisher) in serum-free Opti-MEM for 6 h at 37°C. After treatment, the medium was replaced with fresh complete medium, and cells were incubated for another 24 h. For CCK-8 analysis, 10 μL of CCK-8 reagent was added to each well, followed by incubation at 37°C for 2 h. Absorbance at 450 nm (reference: 650 nm) was measured using a Varioskan™ LUX microplate reader (Thermo Fisher). Untreated cells served as the 100% viability control, and blank wells (medium only) were used for background correction.

### Gastric cancer organoid culture and mRNA transfection

Gastric Cancer (GC) organoids were derived from peripheral blood and tissue samples of GC patients undergoing surgical resection at the Second Affiliated Hospital of Army Medical University, with approval from the Medical Ethics Committee (Protocol ID: 2022-096-01). All procedures were in accordance with the Declaration of Helsinki.

Organoid Culture: Organoids were suspended in cell recovery solution (BD Falcon, 354253) and incubated for 30 minutes on ice with constant shaking (100 rpm), followed by centrifugation (200 g, 5 minutes). Organoids were digested with TryPLE Express (Gibco, 12605-028) and plated at 2,000 cells per well in 6-well plates coated with 20 µL Matrigel (Corning, 356231). They were cultured in GC organoid basal medium (bioGenous, K2179-GC) at 37°C in a 5% CO2 incubator, with medium changes every other day. Organoids were passaged between day 7 and day 10, and experiments were performed from passages 7 to 10. mRNA Transfection: Organoids were suspended in cryopreservation medium (bioGenous, E238023) and incubated for 30 minutes on ice with shaking (100 rpm), followed by centrifugation (200 g, 5 minutes). The organoids were plated in 24-well plates pre-treated with anti-adherence solution (bioGenous, E238002) at 200 organoids per well in 100 µL basal medium. LNP/Glu-LNP was mixed with organoid basal medium at a 1:200 (w/v) ratio of IVT mRNA to medium. A total of 1 µg IVT mRNA was added per well in triplicates. After 12 h of incubation at 37°C in a 5% CO2 incubator, organoids were re-coated with Matrigel. They were incubated for another 24 h before assessing eGFP expression via confocal microscopy (Olympus FV3000) and flow cytometry (BD FACSCanto II). Luciferase activity was measured using a microplate reader (Thermo Fisher Scientific Varioskan Lux).

### Animal studies

Ethical Compliance and Experimental Design The animal research protocol was conducted in compliance with international laboratory animal care standards (Approval No. AMUWEC20223412) and authorized by the Institutional Animal Ethics Committee of TMMU. The experimental design incorporated four representative rodent models: 1) female BALB/c mice (6-8 weeks); 2) C57BL/6 mice (6-8 weeks); 3) SD rats (8-10 weeks); 4) Syrian hamsters (7-8 weeks), all commercially acquired from Vital River Laboratories (Beijing, China).

Housing and Pre-experimental Management All subjects were housed in individually ventilated cages (IVC systems) under strict SPF conditions, maintaining controlled environment parameters: temperature 22±1°C, humidity 55±5%, with 12 h circadian rhythm regulation. Sterilized feed (autoclaved pellet diet) and acidified water (pH 2.5-3.0) were provided ad libitum. Following a 7-day acclimatization period involving daily health monitoring (body weight, coat condition, activity levels), animals underwent stratified randomization based on weight-matching principles prior to experimental interventions.

### In vivo bioluminescence and In vivo biodistribution

Animals (BALB/c, C57BL/6 mice, SD rats, and Syrian hamsters) were intranasally administered with LNP/Glu-LNP formulations containing 2 μg (mice) or 10 μg (rats/hamsters) of Fluc-mRNA for bioluminescence tracking, while a parallel cohort received DiD-labelled nanoparticles (2 μg mRNA equivalent) for biodistribution analysis. At 6 h post-administration, all subjects were anesthetized with 2% isoflurane and injected intraperitoneally with D-luciferin (150 mg/kg). Whole-body bioluminescence imaging was performed using an IVIS Lumina III system (60-120 s exposure, FOV 15-25 cm), followed by immediate euthanasia and collection of major organs (brain, lungs, heart, liver, spleen, kidneys). Excised organs were subjected to fluorescence imaging (Ex/Em: 640/680 nm) without perfusion, with bioluminescence signals quantified via Living Image 4.5 software and fluorescence intensities normalized to tissue mass.

### Histopathological analysis

At 6 h post-intranasal administration of LNP or Glu-LNP (2 μg for mice), major organs (lungs, brain, intestines, liver, spleen, kidney) were harvested, fixed in 4% paraformaldehyde, and processed through paraffin embedding. Tissue sections (5 μm thickness) were stained with hematoxylin and eosin (H&E) and analyzed using a Nikon Eclipse Ci-L microscope to assess pathological changes.

### Cytokine detection in lung homogenates, BALF and serum

Cytokine levels (TNF-α, IL-6, IL-1β) in lung homogenates, BALF, and serum were quantified post-intranasal administration of LNP/Glu-LNP using ELISA kits (Dakewe Biotech: TNF-α #1217203, IL-6 #1210602, IL-1β #1210112). Samples were diluted serially and transferred to pre-coated plates, incubated at 37°C for 90 min. After washing, HRP-conjugated streptavidin was added (37°C, 30 min), followed by TMB substrate (Abcam #ab171522; 5 min incubation). Reactions were terminated with stop solution (Beyotime #P0215), and absorbance was measured at 450 nm using a Varioskan lux microplate reader (Thermo Fisher).

### Systemic biochemical safety evaluation

To assess the short-term systemic safety of Glu-LNP, BALB/c mice were intranasally administered with the test formulations (50 μL/mouse). At 6 h post-administration, whole blood was collected via retro-orbital sampling and allowed to clot at room temperature for serum separation. The obtained serum samples were submitted to Servicebio (Wuhan, China) for comprehensive biochemical analysis. Biochemical parameters included liver function markers-alanine aminotransferase (ALT), aspartate aminotransferase (AST), and alkaline phosphatase (ALP); renal function markers-creatinine (CREA) and blood urea nitrogen (BUN); and metabolic indicators—glucose (GLU) and glycated serum protein (GSP). All measurements were conducted following standard clinical chemistry protocols using an automated biochemical analyzer.

### Transcriptomic Analysis

Lung tissues were collected from Balb/c mice (n=3 per group) 6 h after administration of PBS, Glu-LNP, or LNP formulations. Tissues were fixed in 4% paraformaldehyde (PFA), and total RNA was isolated using the RNeasy Mini Kit (Qiagen) following DNase I treatment to eliminate genomic DNA contamination. RNA quality was assessed using a Bioanalyzer 2100 (Agilent Technologies), and samples with RNA Integrity Number (RIN) ≥8.0 were selected for library preparation. Stranded mRNA sequencing libraries were constructed with the NEBNext Ultra II RNA Library Prep Kit (Illumina) and sequenced on an Illumina NovaSeq 6000 platform (paired-end 150 bp) by Majorbio Bio-pharm Technology Co., Ltd (Chongqing, China). Raw sequencing data were processed on the Majorbio Cloud Platform (https://cloud.majorbio.com) using the following pipeline: (1) Quality control via Fastp v0.23.2 to trim adapters and filter low-quality reads (Q-score <20); (2) Alignment to the GRCm39 mouse reference genome using STAR v2.7.10a with default parameters; (3) Quantification of gene expression levels via featureCounts v2.0.3; (4) Differential expression analysis using DESeq2 v1.38.3 (adjusted p-value <0.05, |log2 fold change| >1); (5) Functional enrichment analysis of DEGs through KEGG and GO databases (Fisher’s exact test, FDR-corrected p-value <0.1).

### Murine models of lung cancer and therapeutic evaluation

Metastatic lung cancer model: C57BL/6 mice were intravenously injected with 1×10⁶ B16-F10-Luc cells suspended in 200 μL PBS on day 0 to establish a lung metastatic model. Mice were randomized into three groups (LNP, Glu-LNP, PBS; n=9/group). On days 6, 9, and 12, IL-12 mRNA (2 μg/mouse) encapsulated in LNP or Glu-LNP was administered via inhalation delivery, while the PBS group received phosphate-buffered saline. Tumor progression was monitored at 72 h intervals from day 9 to day 21 via bioluminescence imaging (IVIS Spectrum, PerkinElmer) after intraperitoneal injection of 150 mg/kg D-luciferin potassium salt. Lungs were excised, weighed to quantify metastatic burden, and visually inspected for gross lesions.

Orthotopic lung cancer model: LLC-Luc (Lewis lung carcinoma) cells were used to establish a primary lung tumor model. C57BL/6 mice were anesthetized and intratracheally instilled with 5×10⁵ LLC cells suspended in 50 μL PBS under sterile conditions. Mice were randomized into three groups (LNP, Glu-LNP, PBS; n= 9/group). On days 6, 9, and 12, IL-12 mRNA (2 μg/mouse) encapsulated in LNP or Glu-LNP was administered via inhalation delivery, while the PBS group received phosphate-buffered saline. Tumor progression was monitored at 72 h intervals from day 9 to day 21 via bioluminescence imaging (IVIS Spectrum, PerkinElmer) after intraperitoneal injection of 150 mg/kg D-luciferin potassium salt. On day 21, lungs were harvested, weighed, and fixed in 4% paraformaldehyde for histological evaluation.

### RT-qPCR Analysis of Tyrp1 Expression in Lung Tissue

Total RNA was extracted from mouse lung tissues using the OMEGA MicroElute Total RNA Kit (R673101). One microgram of RNA was reverse-transcribed into cDNA using the Takara PrimeScript RT Reagent Kit (RR037B). Quantitative PCR was conducted using the YEASEN Hieff SYBR Green Master Mix Kit (11201ES08) with gene-specific primers.

### Intracellular ion detection

Chloride Ion Detection: At 6 h post-LNP/Glu-LNP administration, 16HBE cells were washed three times with Krebs-HEPES buffer (2 min per wash). After removing residual buffer, 1 mL of MQAE working solution (MedChemExpress, HY-D0090) was added and incubated at room temperature for 30–60 minutes. Cells were washed twice with PBS (5 min per wash), and chloride levels were visualized using fluorescence microscopy with excitation/emission wavelengths of 350/460 nm.

Calcium Ion Detection: At 6 h post-LNP/Glu-LNP administration, 16HBE cells were washed three times with calcium/magnesium-free HBSS (Servicebio, G4203-500ML). Subsequently, 0.5 μL Fluo-4 AM (1 mM stock, MedChemExpress, HY-101896) was added to achieve a final concentration of 5 μM and incubated at 37°C for 1 h. Cells were washed twice with HBSS (5 min per wash), and intracellular calcium levels were analyzed using fluorescence microscopy with excitation/emission wavelengths of 494/516 nm.

### Western blot

Total protein was extracted from lung tissues using RIPA lysis buffer containing protease and phosphatase inhibitors (Servicebio, G2002). Protein concentration was quantified using a BCA assay kit (Servicebio, G2026). Equal amounts of denatured protein were resolved by SDS-PAGE and transferred onto PVDF membranes (0.45 µm, Servicebio, G6015). Membranes were blocked in 5% skim milk and incubated overnight at 4 °C with primary antibodies against the target protein. After chemiluminescent detection (ECL, Servicebio, G2161) and imaging (Servicebio SCG-W2000), the membranes were incubated with enhanced antibody stripping buffer (Servicebio, G2079) to remove the previously bound antibodies. This procedure was repeated to sequentially detect additional target proteins on the same membrane. Internal control antibodies (β-actin) were applied at the final stage and visualized using the same procedure. All grayscale quantification was performed using AIWBwell™ analysis software (Servicebio), and relative expression levels were calculated as the ratio of target protein signal to internal control.

### Statistical Analysis

Unless otherwise specified, data are presented as mean ± SEM. Statistical analyses were conducted using Prism 8 (GraphPad Software). Pairwise comparisons were evaluated by Welch’s t-test, while multi-group comparisons were analyzed via ANOVA with post-hoc tests. All tests were two-tailed, and significance was defined as P < 0.05.

## Acknowledgements

1. S. Guan acknowledge support for the research described in this study from the National Natural Science Foundation of China (NSFC, Grant No. 32370993 and 82173764), and the major project of Study on Pathogenesis and Epidemic Prevention Technology System (2021YFC2302500 and 2024YFC2310804) by the Ministry of Science and Technology of China. We thank Dr. X. Zhu (Shanghai Vitalgen BioPharma Co., Ltd) for helpful discussion. L. Xu acknowledges the support of the China Scholarship Council (CSC) (202508500053).

## Author contributions

S.G. conceived and directed the project. S.G. supervised the research and contributed experimental materials. L.X, Y.C., R.L., F.Y., M.L and Q.X. performed experiments. L.X., C.L. and S.G. completed the design of the machine learning model and finalized the curation of sugar-related datasets. L.X., Y.C., Q.X. and C.L. analyzed data. T.C. and C.L. was responsible for the bioinformatics and RNA sequencing data analysis. S.G., L.X., C.L., and T.C. wrote the manuscripts with help and comments from all authors.

## Data availability

All data supporting the results in this study are available within the paper and its Supplementary Information. Source data are provided with this paper.

## Code availability

The source code used for the machine learning framework is available from the corresponding author upon reasonable request.

## Extended Data Figures

**Extended Data Fig. 1.**
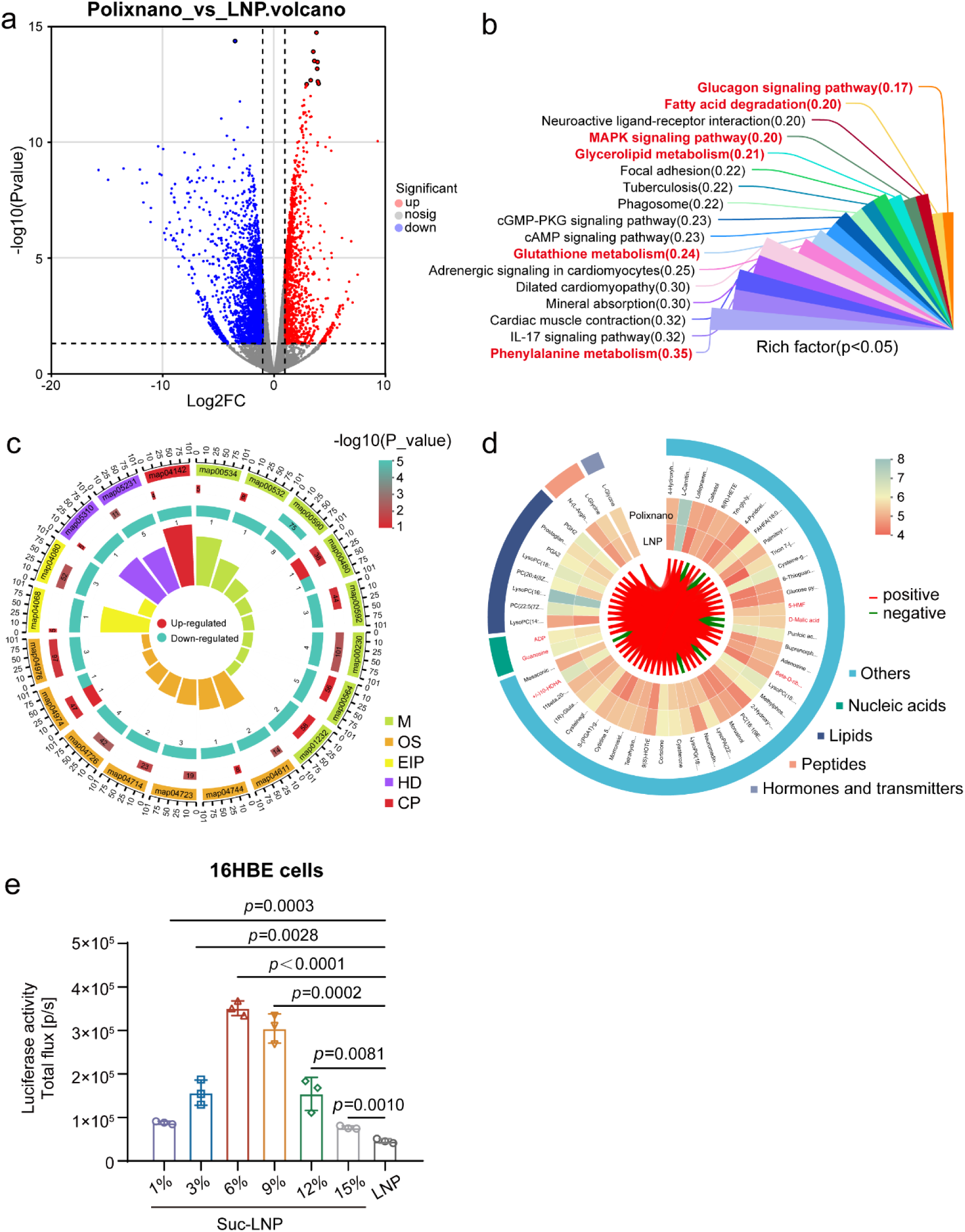
Transcriptomic and Metabolomic Profiling of Lung Tissues Following PolixNano and LNP Treatment. **a**, Volcano plot of differentially expressed genes between PolixNano-treated and LNP-treated lung tissues, highlighting significantly upregulated (red) and downregulated (blue) transcripts. **b,** KEGG pathway enrichment analysis of differentially expressed genes identified in lung tissues following PolixNano treatment compared to LNP, showing the top 20 enriched pathways ranked by rich factor (p < 0.05). Four metabolism-related pathways are marked in red, including glycerolipid metabolism, fatty acid degradation, glutathione metabolism, and phenylalanine metabolism. **c,** KEGG enrichment analysis of significantly altered metabolites identified through untargeted metabolomics comparing PolixNano and LNP treatments. Red and green bars represent upregulated and downregulated metabolites, respectively. Functional categories of the enriched pathways are color-coded as follows: M (Metabolism), OS (Organismal Systems), EIP (Environmental Information Processing), HD (Human Diseases), and CP (Cellular Processes). **d,** Chord diagram showing the correlation between significantly altered metabolites and enriched KEGG pathways, with red highlighting metabolites associated with carbohydrate metabolism, including glucose pyruvate, D-malic acid, beta-D-ribofuranose, and 5-hydroxymethylfurfural, etc.. **e,** mFluc expression in 16HBE cells transfected with Suc-LNP at varying concentrations (1%-15% w/v), showing enhanced expression compared to the LNP counterpart (n=3 biologically independent samples).

**Extended Data Fig. 2.**
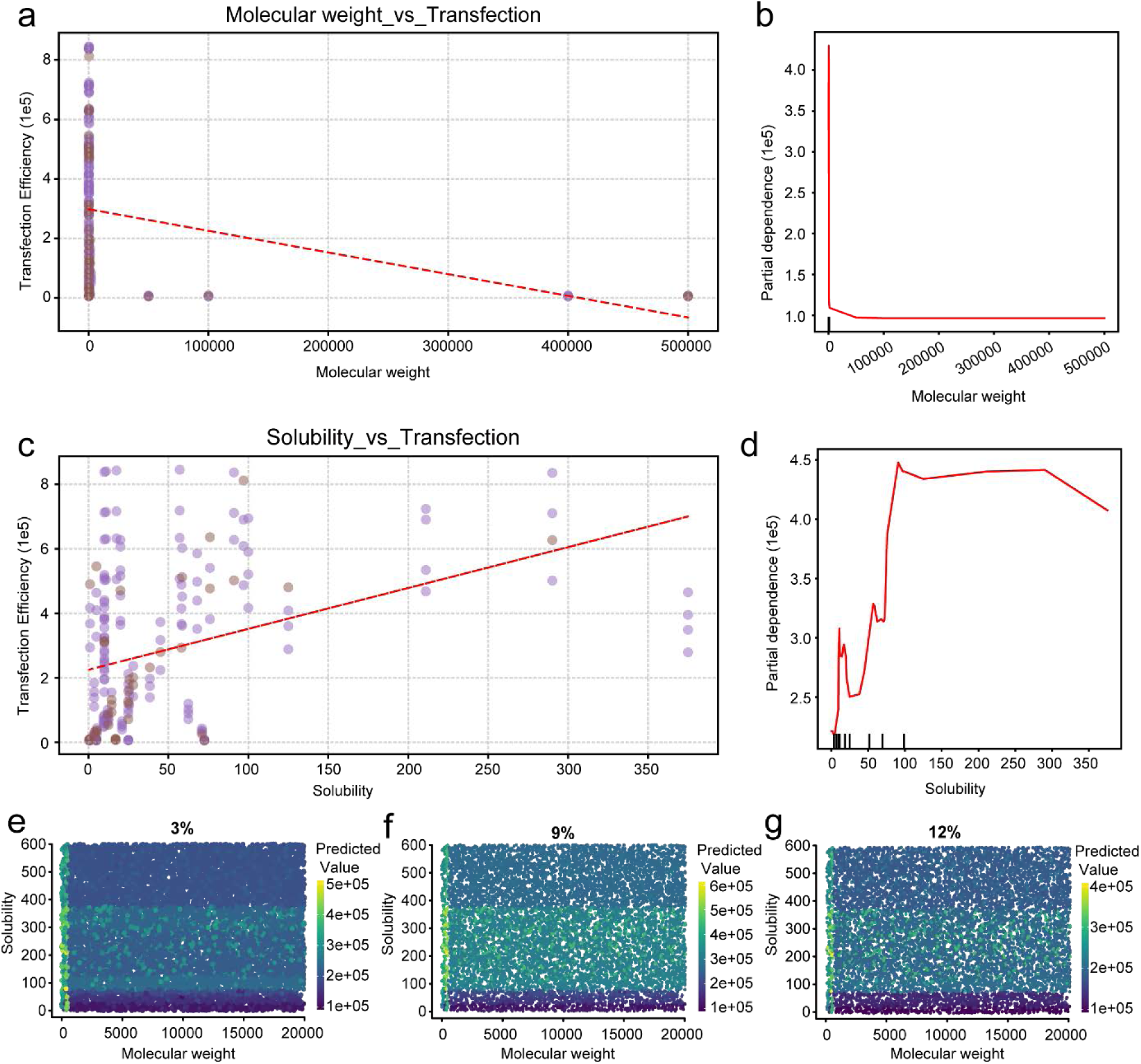
Machine Learning–Driven Identification of Sugar Characteristics Influencing LNP Transfection Efficiency. **a-b**, Correlation analysis between molecular weight and transfection efficiency (a) and partial dependence of molecular weight(b) derived from XGBoost modeling based on the experimental dataset. Each data point represents an individual sugar-LNP formulation. **c-d,** Correlation analysis between solubility and transfection efficiency (c) and partial dependence of solubility (d) derived from XGBoost modeling. **e-g,** Predicted mFluc transfection efficiency across different molecular weights and solubility ranges, as determined by virtual screening using the XGBoost model at varying sugar concentrations (3%, 9%, and 12% w/v).

**Extended Data Fig. 3.**
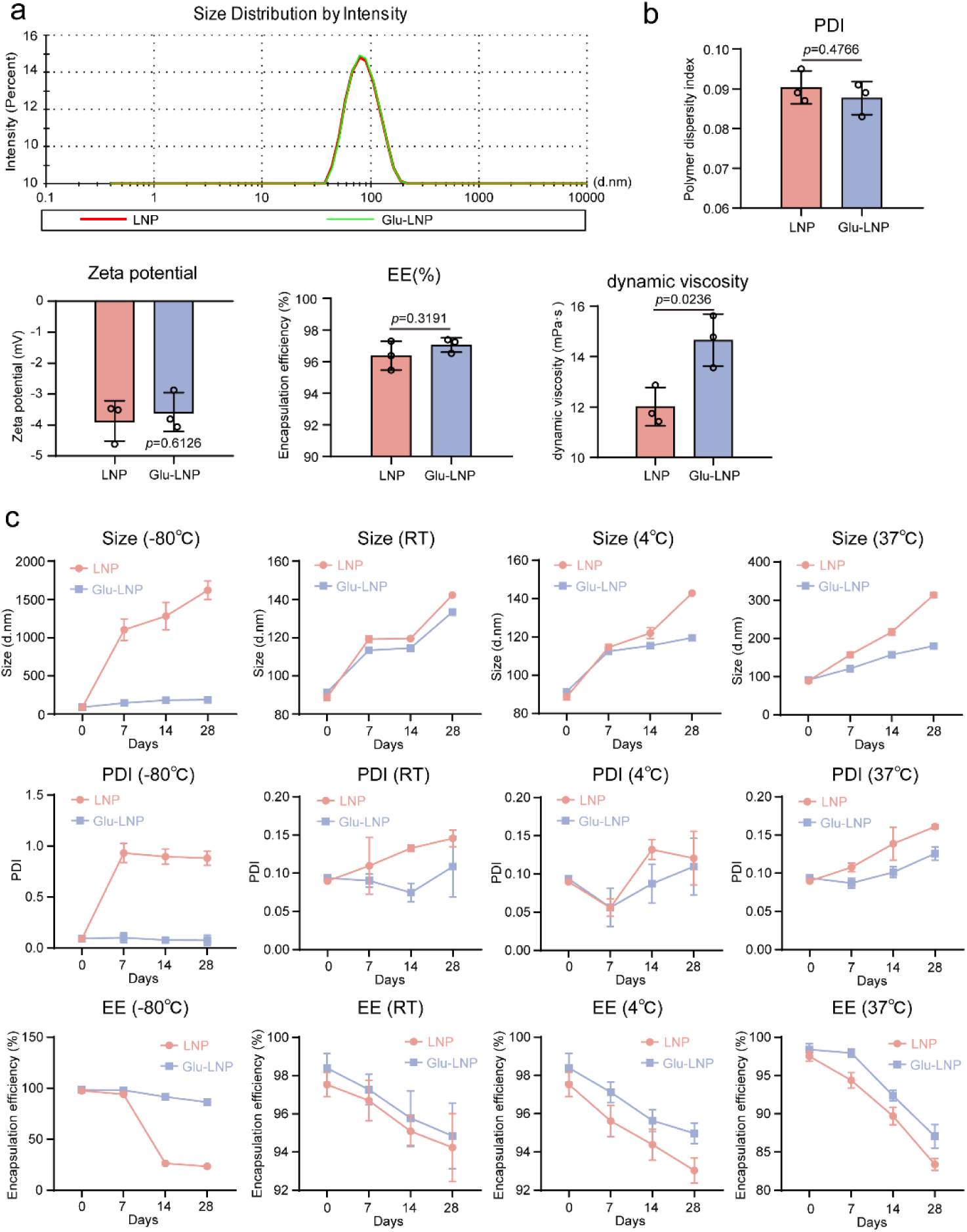
Physicochemical Characterization and Stability Assessment of Glu-LNP Versus Conventional LNP Formulations. **a**, Dynamic light scattering (DLS) analysis comparing particle size distribution profiles of Glu-LNP and LNP formulations. **b,** Comparison of key physicochemical properties between Glu-LNP and LNP formulations, including polydispersity index (PDI), zeta potential, mRNA encapsulation efficiency (EE), and dynamic viscosity (n=3 biologically independent samples). **c,** Stability evaluation of Glu-LNP and LNP formulations under different storage temperatures (−80 °C, 4 °C, room temperature, and 37 °C), comparing changes in particle size, PDI, and EE over time.

**Extended Data Fig. 4.**
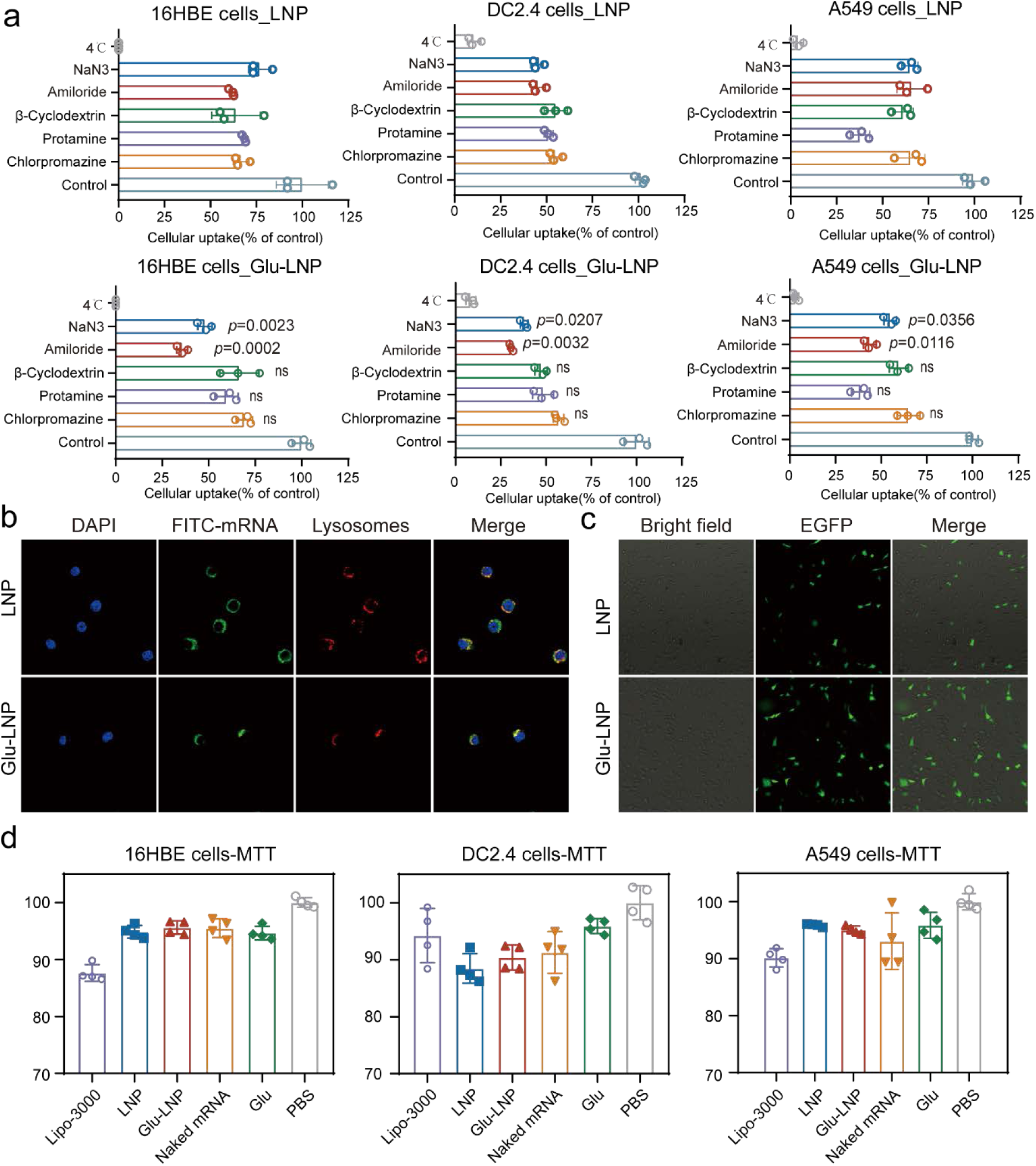
Additional mechanistic and biosafety evaluation supporting glucose-mediated enhancement of transfection. **a**, Cellular uptake inhibition profiles of mFITC encapsulated in standard LNP in 16HBE, DC2.4, and A549 cells (n=3 biologically independent samples). **b,** Confocal microscopy images confirming lysosomal escape of mFITC encapsulated in Glu-LNP or LNP in DC2.4 cells. Channels represent mFITC (green), lysosomes (red), nuclei (blue). Scale bars: 15 µm. **c,** Representative fluorescence microscopy images of DC2.4 cells transfected with mEGFP encapsulated in Glu-LNP or LNP after 6 h incubation. Scale bar: 100 µm. **d,** Cell viability was assessed using the methyl thiazolyl tetrazolium (MTT) assay in DC2.4, 16HBE, and A549 cells after treatment with Lipofectamine 3000, LNP, Glu-LNP, naked mRNA, glucose, or PBS as a negative control (n=6 biologically independent samples).

**Extended Data Fig. 5.**
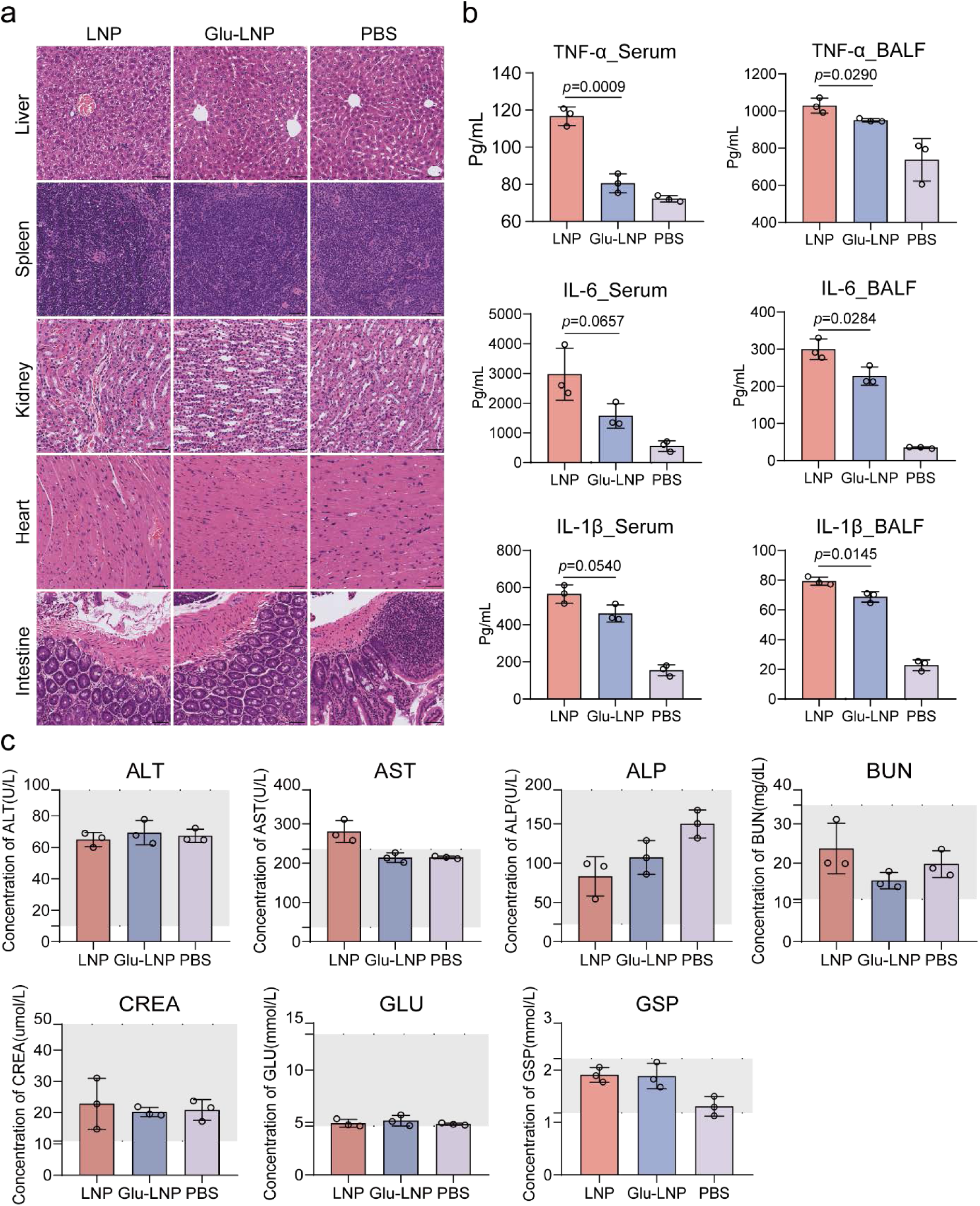
Additional in vivo evaluations of Glu-LNP. **a**, H&E staining of heart and intestine sections collected from mice at 6 h post-intranasal instillation with Glu-LNP or LNP. Scale bar: 50 μm. **b,** Corresponding levels of TNF-α, IL-6, and IL-1β in serum and BALF collected from separately treated mice 6 h post-administration of PBS, LNP, or Glu-LNP (n=3 biologically independent samples). Cytokine concentrations were measured by ELISA. **c,** Complete blood chemistry analysis of mice treated with intranasally administered Glu-LNP (n=3 biologically independent samples). Grey regions indicate normal reference ranges. ALT: alanine aminotransferase; AST: aspartate aminotransferase; ALP: alkaline phosphatase; BUN: blood urea nitrogen; CREA: creatinine; GLU: glucose; GSP: glycated serum protein.

**Extended Data Fig. 6.**
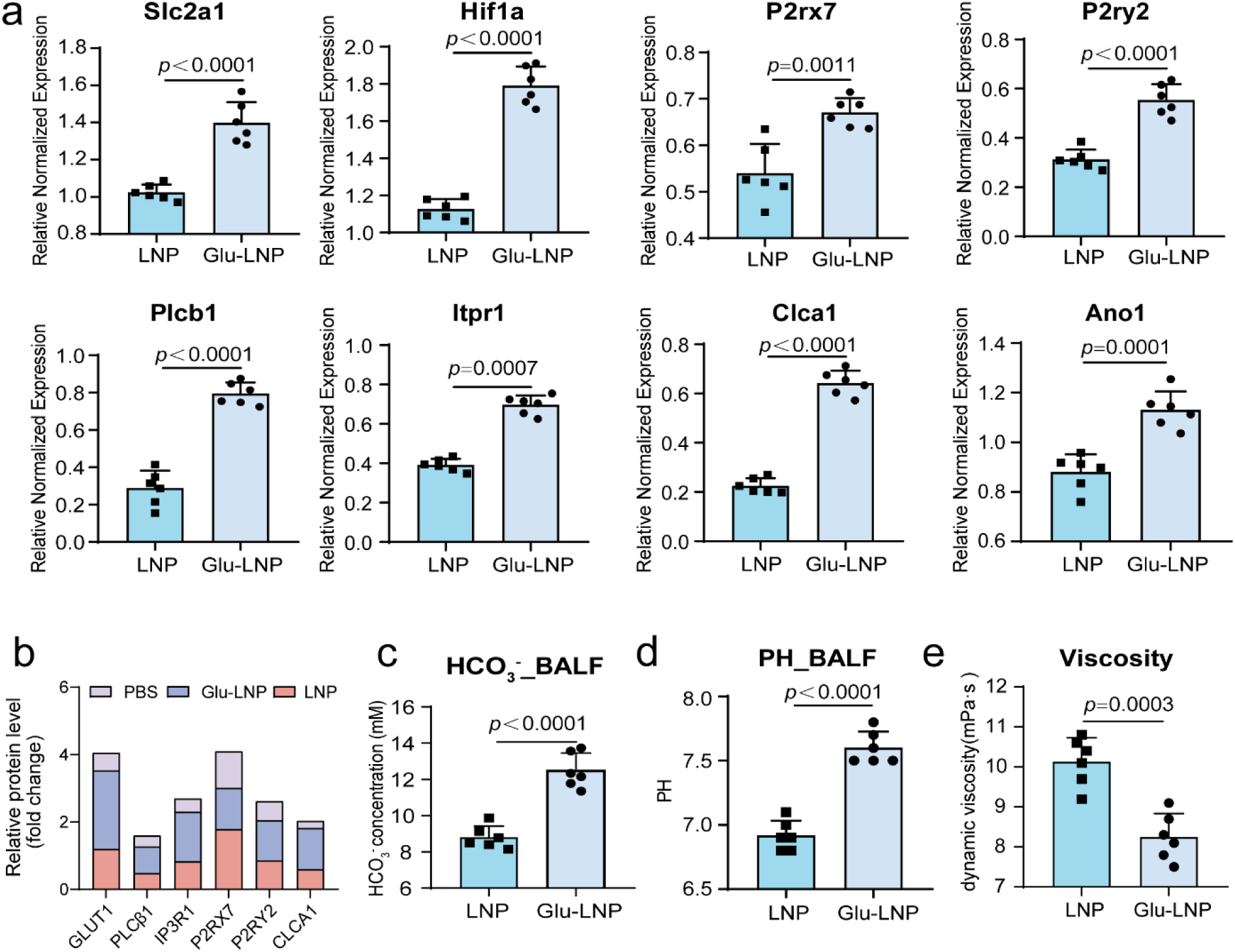
Glu-LNP Activates the P2Y2–IP₃–CLCA1 Axis to Modulate Airway Ion Transport and Mucus Properties. **a**, Differential gene expression analysis showing key genes involved in the P2Y2-IP₃-CLCA1 signalling axis in mouse lung tissues treated with Glu-LNP or LNP, measured by RT-qPCR (n=6 biologically independent samples). **b,** Quantification of protein expression levels of GLUT1, PLCβ1, P2RX7, P2RY2, IP3R1 and CLCA1 by grayscale value analysis of Western blot results. **c-d,** Quantification of pH value and HCO₃⁻ concentration in BALF from Glu-LNP and LNP treated mice (n=6 biologically independent animals). **e,** Quantification of mucus viscosity in simulated airway mucus samples following 6 h incubation with NaHCO₃ or PBS control (n=6 biologically independent samples).

## Supplementary information

### SUPPLEMENTARY TABLES

**Supplementary Table 1.**
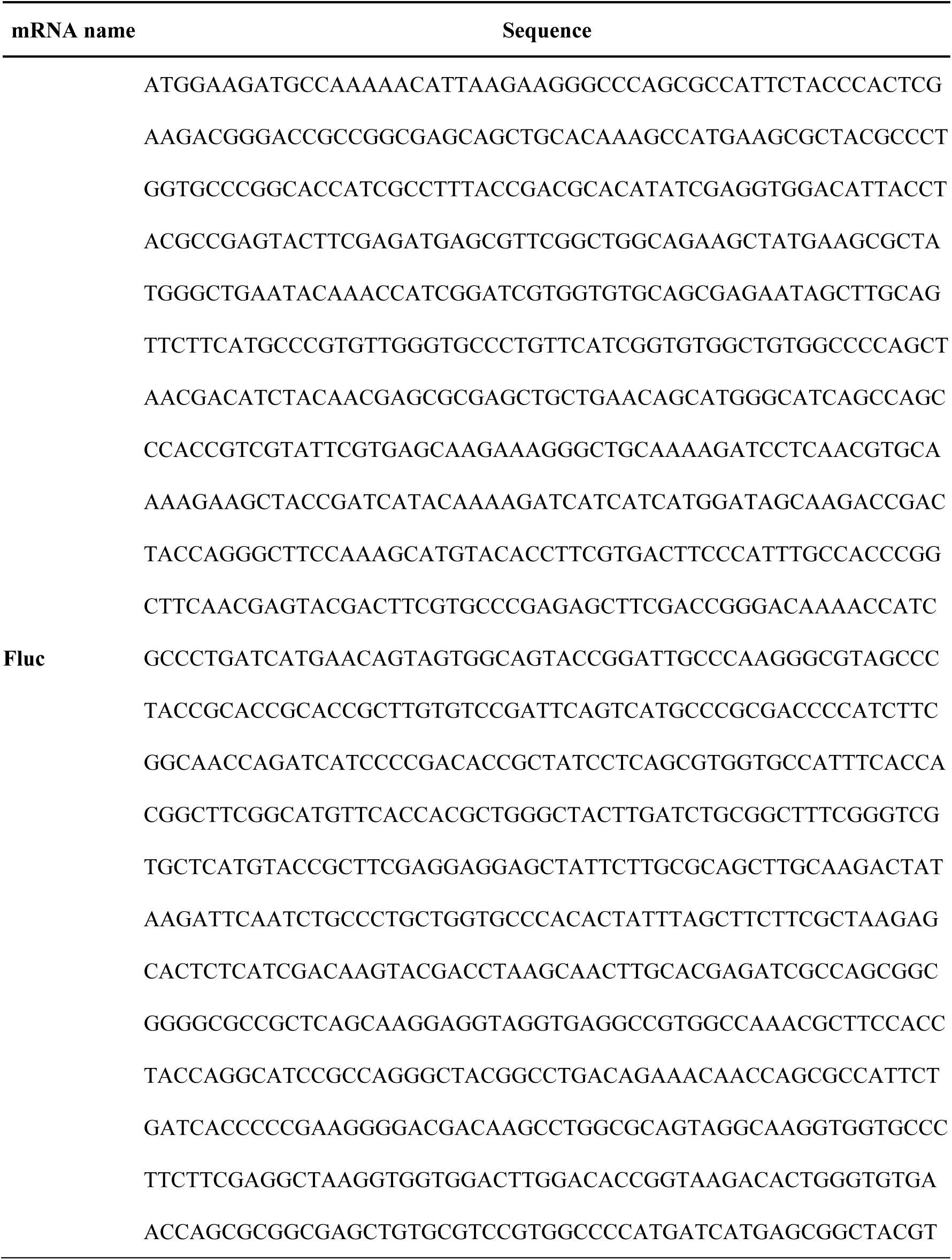
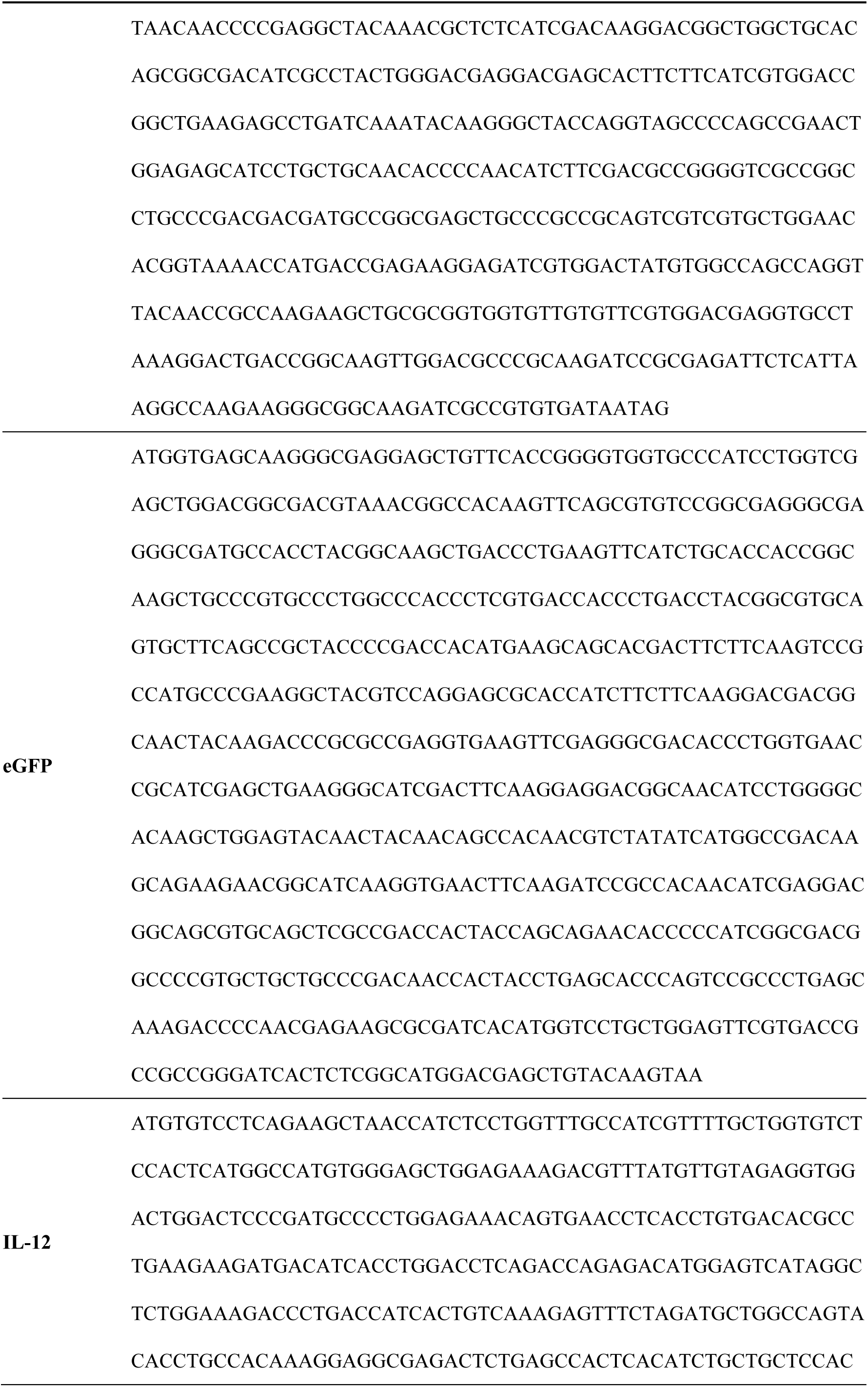
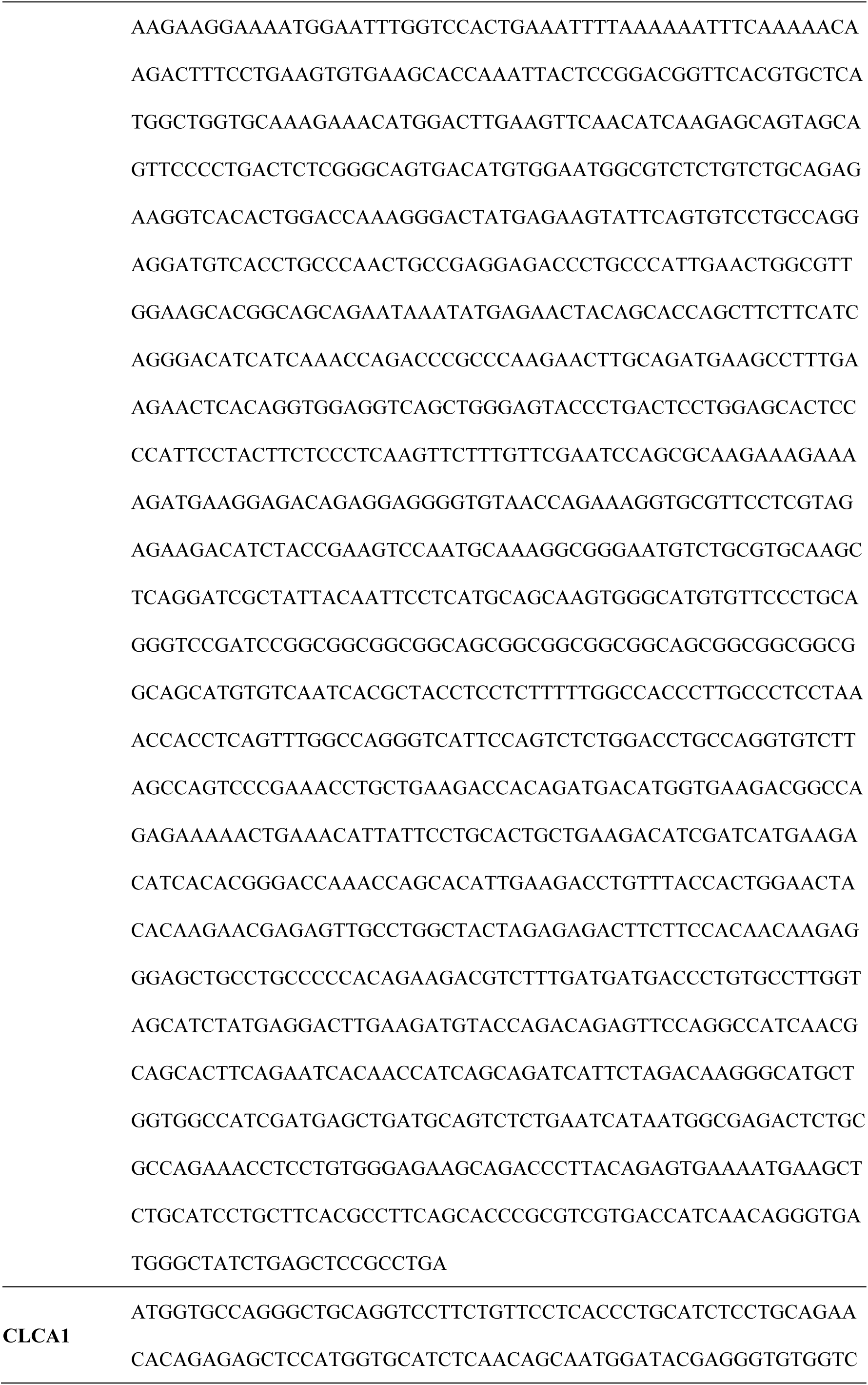
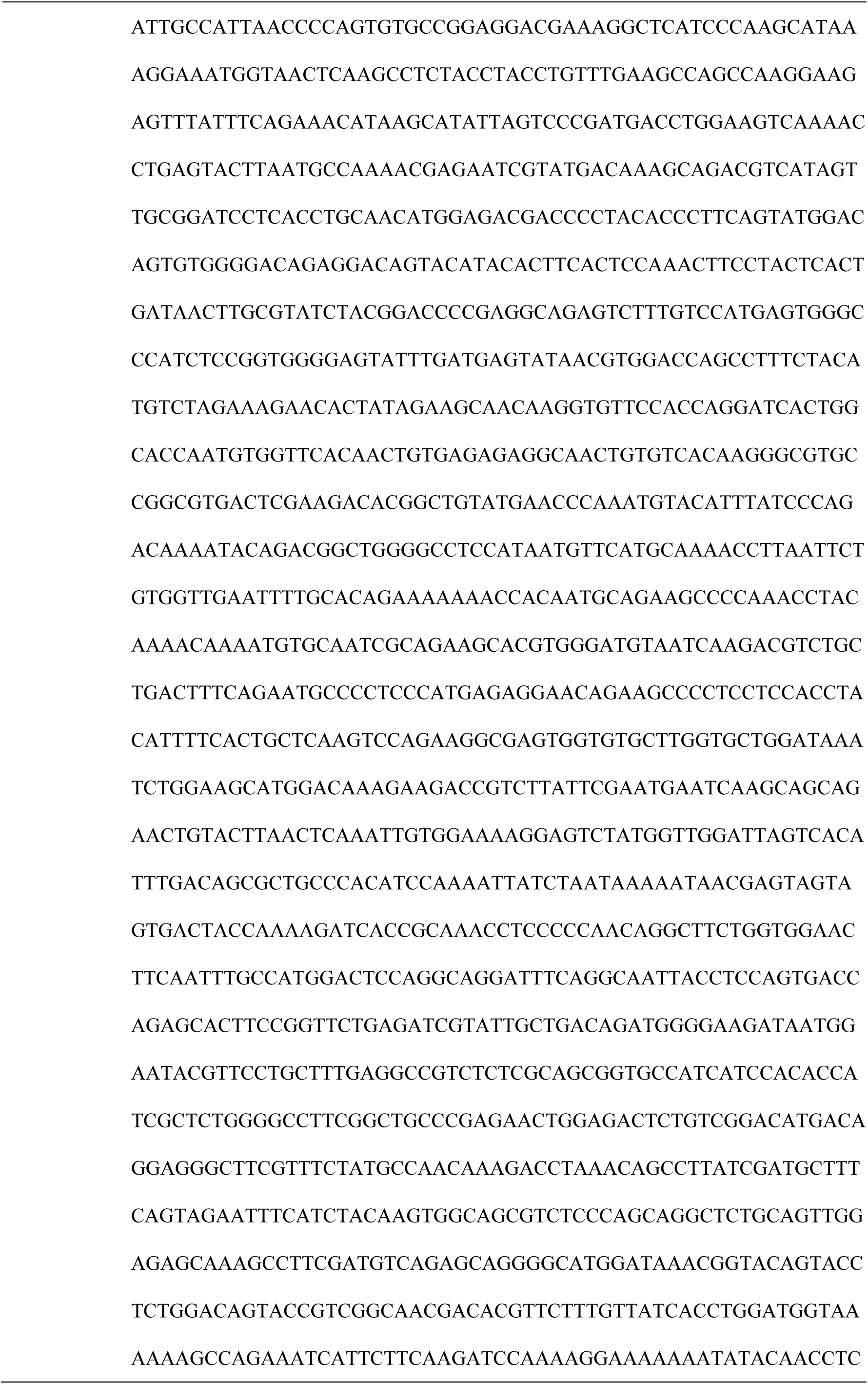
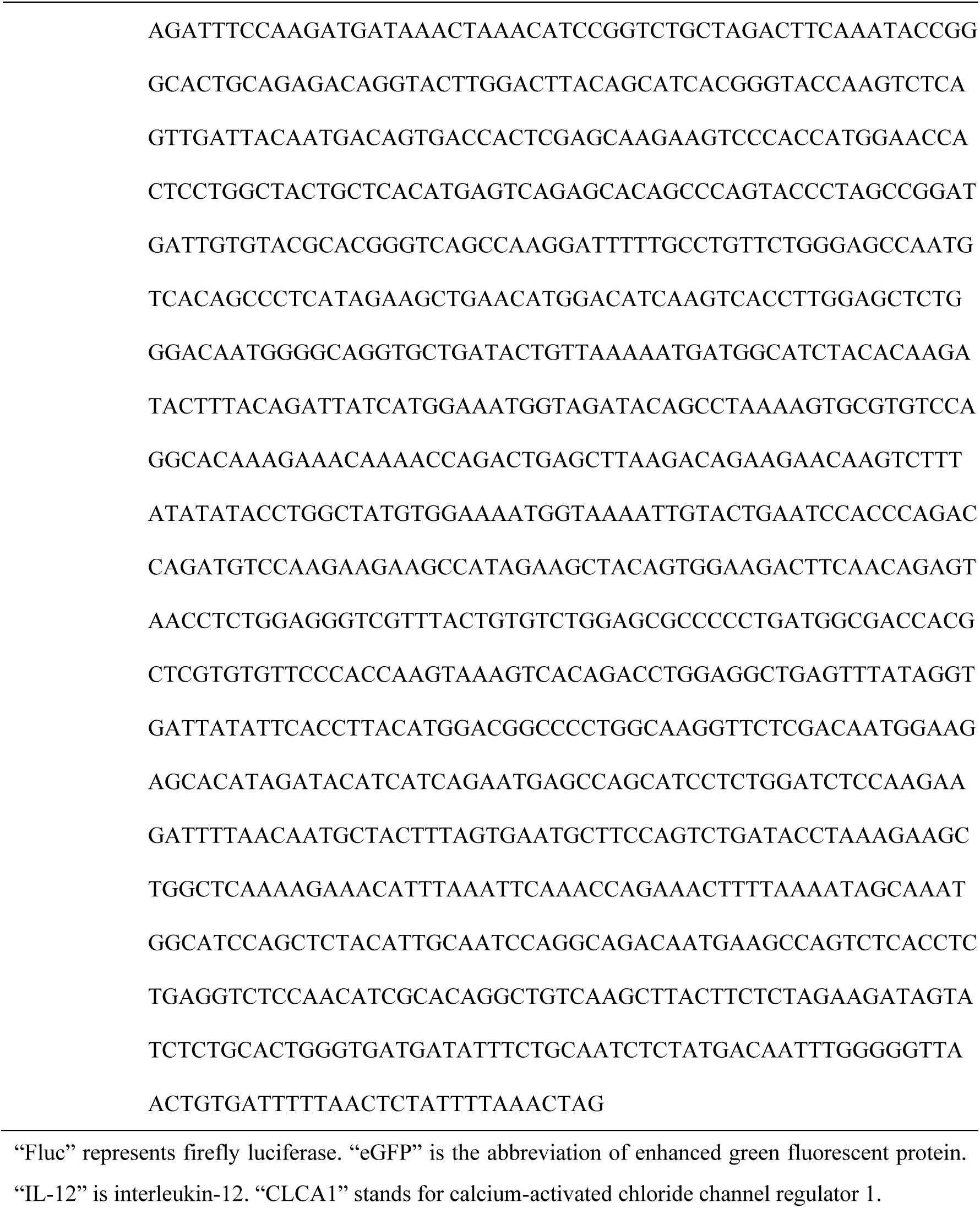
The open reading frame (ORF) sequences of in vitro transcribed mRNA adopted in this study.

**Supplementary Table 2.**
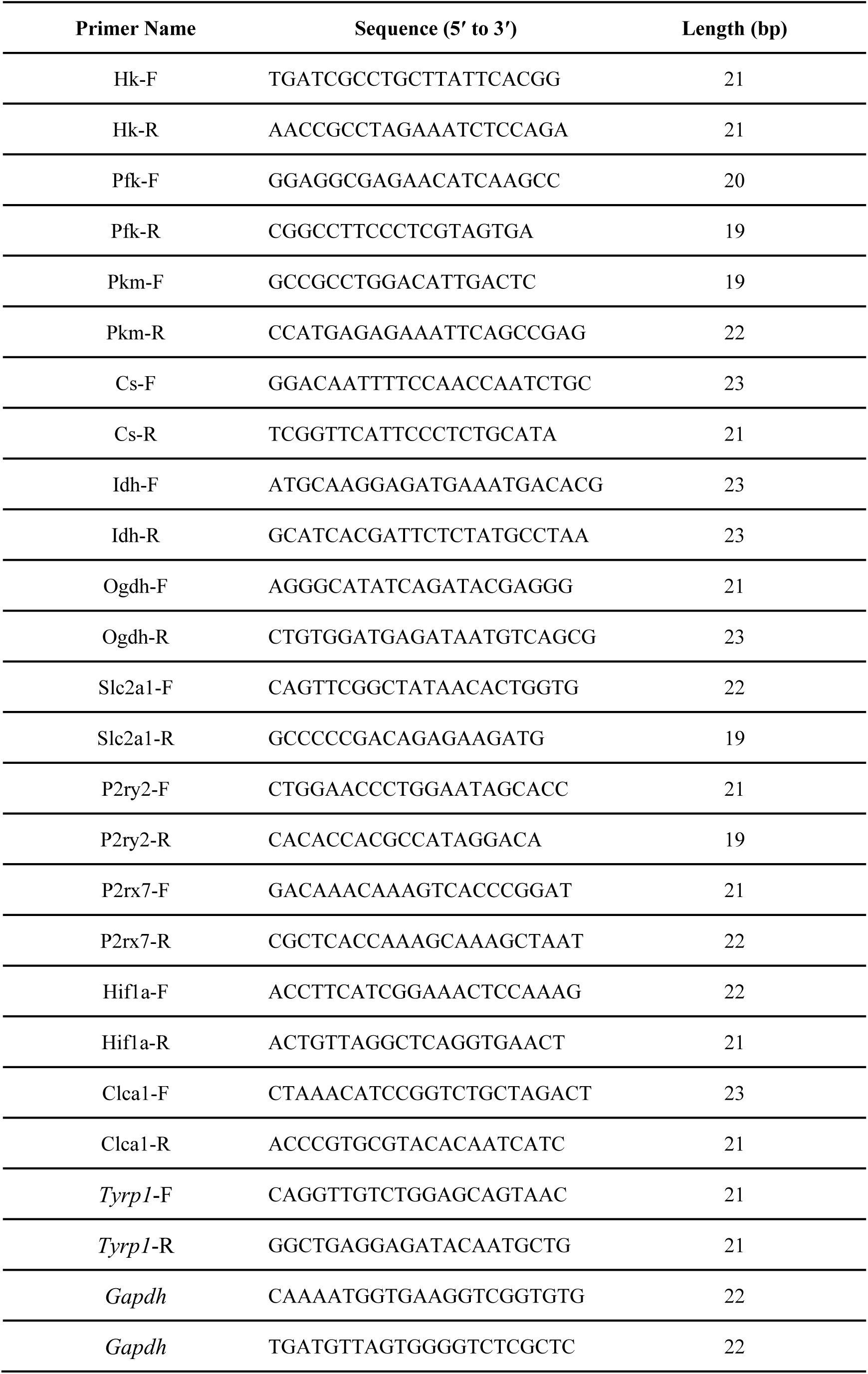
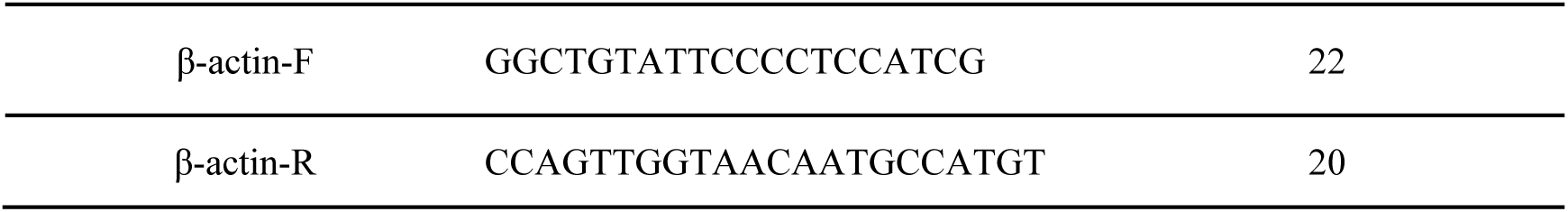
Primer sequences used for RT-qPCR.

